# Enhancer prediction in the human genome by probabilistic modelling of the chromatin feature patterns

**DOI:** 10.1101/804625

**Authors:** Maria Osmala, Harri Lähdesmäki

## Abstract

**Background:** The binding sites of transcription factors (TFs) and the localisation of histone modifications in the human genome can be quantified by the chromatin immunoprecipitation assay coupled with next-generation sequencing (ChIP-seq). The resulting chromatin feature data has been successfully adopted for genome-wide enhancer identification by several unsupervised and supervised machine learning methods. However, the current methods predict different numbers and different sets of enhancers for the same cell type and do not utilise the pattern of the ChIP-seq coverage profiles efficiently.

**Results:** In this work, we propose a PRobabilistic Enhancer PRedictIoN Tool (PREPRINT) that assumes characteristic coverage patterns of chromatin features at enhancers and employs a statistical model to account for their variability. PREPRINT defines probabilistic distance measures to quantify the similarity of the genomic query regions and the characteristic coverage patterns. The probabilistic scores of the enhancer and non-enhancer samples are utilised to train a kernel-based classifier. The performance of the method is demonstrated on ENCODE data for two cell lines. The predicted enhancers are computationally validated based on the transcriptional regulatory protein binding sites and compared to the predictions obtained by state-of-the-art methods.

**Conclusion:** PREPRINT performs favorably to the state-of-the-art methods, especially when requiring the methods to predict a larger set of enhancers. PREPRINT generalises successfully to data from cell type not utilised for training, and often the PREPRINT performs better than the previous methods. The PREPRINT enhancers are less sensitive to the choice of prediction threshold. PREPRINT identifies biologically validated enhancers not predicted by the competing methods. The enhancers predicted by PREPRINT can aid the genome interpretation in functional genomics and clinical studies.

**Availability:** https://github.com/MariaOsmala/preprint

## Background

In recent years, there have been many papers describing computational methods to predict genomic regulatory enhancers. The methods have adopted the chromatin feature data produced by the next-generation sequencing technologies. Developing methods to identify enhancers in the human genome is important, as enhancers are estimated to be over-represented (60–80%) in the discoveries of genome-wide association studies (GWAS) aiming to detect single nucleotide polymorphisms (SNPs) associated with both rare and common diseases [1, 2, 3]. Moreover, enhancers have been ascertained to be the main regulators of the cell type-specific gene expression, and they have an important function in cell differentiation [4, 5, 6]. Therefore, the accurate identification of enhancers could support the interpretation of the GWAS findings, the results from the functional genomics studies and finally the deployment of the precision medicine.

The number of enhancers in the human genome is estimated to be in the hundreds of thousands. Enhancers are difficult to locate as they are independent in position, distance, and orientation with respect to their target genes [7, 8], and lack general sequence specificities. Enhancers have been shown to possess certain molecular and structural chromatin features, which can be utilised to locate them genome-wide. Chromatin resides inside the cell nucleus and consists of DNA wrapped around nucleosomes and other protein structures. Chromatin features comprise, for example, the binding sites of transcriptional regulatory proteins (TRF) and the epigenetic modifications of the nucleosomal histone tails, i.e., histone modifications. The chromatin features can be quantified genome-wide using high-throughput next-generation sequencing (NGS) techniques. In particular, the Chromatin Immunoprecipitation coupled with sequencing (ChIP-seq) can quantify the chromosomal locations for tens to hundreds of individual TRFs and histone modifications [9, 10]. Thus, various combinations of the chromatin features have been adopted in several studies to locate enhancers [11, 12, 13, 14, 15, 16, 5, 17, 18].

The NGS techniques quantifying the chromatin features produce a large collection of short sequence reads randomly sampled from the input material, e.g., the cell population. In the analysis of the sequence data, the reads are first aligned back to the reference genome. Secondly, for each base pair (bp) or subsequent non-overlapping genomic locus, a count is defined as the sum of reads aligning to the locus. The read counts along the whole genome or a stretch of DNA are denoted as read coverage signals. The high value of the coverage signal at a given locus compared to the background coverage signal corresponds to the enrichment of the particular chromatin feature at the locus. The genome-wide coverage signals of many individual chromatin features can be processed with machine learning methods to cluster and classify the genomic loci. Consequently, several machine learning methods have been developed for the enhancer prediction task; for earlier reviews and comparisons of the approaches, see [19, 20, 21, 22, 23, 24].

Among the most popular machine learning methods to predict enhancers are a hidden Markov model-based unsupervised method, ChromHMM [25, 26], and a supervised Random Forest-based Enhancer identification from Chromatin States (RFECS) [27]. ChromHMM is restricted to modelling binary data, i.e. the chromatin features are either present or absent depending on a predefined threshold. The choice of the threshold is non-trivial, and due to the binarization, the quantitative information of the ChIP-seq coverage is lost. Moreover, ChromHMM considers the signal in 200 base pair (bp) bins and ignores the characteristic pattern of the coverage signal observed at the regulatory regions. In contrast, RFECS considers the chromatin feature coverage vectors extracted in a 2 kilobase (kb) window centred at the genomic loci of interest, and the window is divided into 20 bins of length 100 bps. Hence, one feature of one locus is a 20-dimensional vector. The vectors are employed to train the RFECS classifier. When training RFECS, at each node in a tree, a subset of chromatin features are randomly selected from the feature set, and the single feature that produces the best separation of classes according to a predetermined criterion is utilised to partition the training data. To reduce the dimension of a feature from 20 to 1 at each node, RFECS applies Fisher Linear Discriminant Analysis, an example of a multi-variate node-splitting technique. The authors of RFECS claim that this approach allows the utilisation of both the coverage pattern and the signal intensity. RFECS was demonstrated to outperform the other supervised methods Chromia [28], CSIANN[29], and ChromaGenSVM [30]. RFECS does not make any distributional assumptions of the data, and the algorithm automatically discovers the optimal subset of the chromatin features for the enhancer prediction task. [27] However, modelling the chromatin feature data according to a distribution that captures the variation of the coverage values has proved to be advantageous [31, 32, 33, 34]. The variation of the coverage values due to the random sampling of the DNA fragments during sequencing can be modelled with the Poisson distribution [31, 32]. However, the coverage data exhibits widespread and consistent overdispersion, i.e., there is a large number of genomic loci with high coverage, much more than expected under a Poisson assumption [35, 36]. The overdispersion results from biological and technical variation, for example, the strength of the interaction between a nucleosome and the DNA, the fragmentation efficiency, the antibody efficiency in the ChIP step, and the local chromatin properties, such as chromatin openness, i.e., DNA accessibility. The observed variation in the read counts among enhancers can also reflect the heterogeneity of the cell line population. Further sources of variability are the ambiguities during the read alignment due to repetitions in the reference genome, and the insufficient sequencing depth. Some of these sources of variation can be taken into account, for example, in ChIP-seq experiments by performing the control experiment without the ChIP step. At a given genomic loci, the variation is usually assumed to originate from a local source.

Despite the vast amount of research, the suggested machine learning methods predict divergent sets of enhancers for the same cell type and do not generalise well between data from separate cell types; in addition, the enhancers predicted by different methods may have unique properties [23, 24, 37, 38]. For example, the lengths of enhancers predicted by the different methods vary (from a few hundreds to a few thousands of bps), and the set of enhancers might not be saturated [38]. The set of enhancers would be saturated if by lowering the prediction thresholds or by analysing more cell types, the collection of enhancers would not change significantly. The inconsistencies between the sets of predicted enhancers likely result from many factors. Firstly, most methods consider the coverage signal in a large genomic window and, hence do not efficiently utilise the pattern of the coverage in subsequent small windows, i.e., bins within the large window. Hence, the methods miss regions with a low coverage which still display the characteristic enhancer patterns. Another challenge involved in enhancer prediction is the calibration of the prediction scores produced by the classifiers, for example, to control the false positive rate. Finally, a crucial challenge in developing a computational enhancer prediction method is the estimation of the accuracy and specificity of the final genome-wide predictions. This is due to a lack of a large gold standard set of human enhancers. At an enhancer, multiple transcription factor-DNA interactions together with the binding of transcription co-factors are required for the enhancer to regulate the target gene expression [39]. As a result, enhancers acquire a high number of co-localised TRF ChIP-seq peaks. Therefore, clusters of TRF ChIP-seq peaks have been employed both to predict enhancers and to validate the predictions [40, 41, 42].

This paper introduces PRobabilistic Enhancer PRedictIoN Tool (PREPRINT), a supervised enhancer prediction method based on probabilistic modelling of the characteristic chromatin feature patterns observed at enhancers and other genomic regions. PREPRINT presumes that the coverage patterns, for example, at individual enhancers resemble each other, although the coverage intensity may vary. Moreover, the coverage values along the characteristic feature patterns are assumed to involve certain dependency structures, such as two-modal peak patterns. In addition, PREPRINT defines two distance measures to quantify “the closeness” or “fit” of the genomic query regions and the characteristic coverage patterns. The distance measure could be as simple as correlation employed in the earlier studies [12, 13]. In contrast, PREPRINT introduces two new probabilistic distance measures. Firstly, PREPRINT assumes Poisson distributed coverage counts with a scaled mean parameter. The scaled mean consists of the characteristic coverage pattern and a scaling parameter. Although an approximation, introducing the characteristic coverage pattern as an estimate of the Poisson mean incorporates the dependency structure of the feature patterns into the model. In addition, the scaling parameter accounts for the coverage signal variability due to the various sources listed above. The scaling parameters are estimated utilising two approaches: either sample-specific scaling parameters are estimated with a maximum likelihood (ML) approach or a global Gamma distribution is estimated for the scaling parameter (Bayesian approach). Furthermore, two probabilistic distance measures are defined depending on the approach employed to learn the scaling parameters. Hence, the two approaches are referred to as PREPRINT ML and PREPRINT Bayesian. PREPRINT is trained and tested on the next-generation sequencing data produced by ENCODE for the myelogenous leukaemia cell line (K562) and the lymphoblastoid cell line (GM12878). PREPRINT predicts genome-wide enhancers in both cell lines. The probability of misclassification for the whole-genome predictions is assessed in advance. This is a requirement for a general tool to predict enhancers on data originating from any cell line. Moreover, the prediction performance of PREPRINT is computationally compared to the state-of-the-art methods RFECS and ChromHMM. Finally, in this work, the enhancers predicted by the computational methods were validated with the largest collection to date of the TRF binding sites produced by the ENCODE consortia [43].

## Results

### Evaluating the classification and generalisation performance of PREPRINT and RFECS

This paper introduces a new enhancer prediction method called PREPRINT. In this section, the classification performances of PREPRINT and the competing method RFECS were compared. The classification performance was evaluated with the area under the receiver operating characteristics curve (AUC). The performance of the methods was evaluated on two small data sets containing chromatin feature data at 1000 enhancers, 1000 promoters, and 2000 random genomic locations. The first data set extracted from the K562 cell line is referred to as training data whereas the second data set extracted from the GM12878 cell line is referred to as test data. The chromatin feature data at the random genomic locations contained data for two separate definitions of the random locations: Firstly, 1000 random locations were sampled uniformly across the whole genome, these locations are referred to as pure random regions. Secondly, another set of 1000 random locations was sampled requiring the chromatin feature signal values at these locations to exceed a certain threshold; hence these locations are referred to as random regions with a signal. Finally, the pure random regions and the random regions with a signal were merged to form the set of combined regions. For more details on the training and test data definition, see the Section Methods.

Next, the training data from the K562 cell line were divided into cross-validation (CV) sets to evaluate the performance of the methods when the training and test data originated from the same cell line (K562). In contrast, the test data from the GM12878 cell line were employed to test the generalisation performance of the methods between the data originating from different cell lines. The AUC values for the different methods and data sets are presented in Table 1. Accordingly, the classification performance of PREPRINT on the K562 CV data set was almost perfect (0.99), and the performance decreased only slightly when predicting enhancers on the GM12878 data with the classifier trained on the K562 data. In addition, the Bayesian version of PREPRINT achieved the best AUC values in both cell lines. For comparison, the methods were also trained and tested on data containing either pure random regions or random regions with a signal. The AUC values from this comparison are presented in Supplementary Table S1, Additional File 1. As expected, the enhancers were easier to separate from the pure random regions than from the random regions with a signal. This was especially the case in the GM12878 cell line. Again, PREPRINT achieved the highest AUC value in all comparisons, PREPRINT Bayesian slightly exceeding the performance of PREPRINT ML in most of the comparisons. When comparing the AUC values of the methods trained on the different random region definitions, the AUC values of methods trained on the combined random regions in Table 1 were between the AUC values of the methods trained on the two random region definitions separately (Supplementary Table S1, Additional File 1). Although the training data definition affected the classification performance on the training and test data sets, the results in the following sections were mainly presented for the methods trained on the combined random regions. To conclude, the classification performance of PREPRINT Bayesian exceeded the performance of PREPRINT ML on data from the same cell line that the methods were trained on (K562). Moreover, PREPRINT achieved superior performance to RFECS when generalising to the data from another cell line (GM12878). However, classifying the small set of highly significant enhancers and promoters is a rather simple task. Thus, next the evaluations of the whole-genome predictions are presented.

**Table 1:** The classification performance (AUC) of PREPRINT and RFECS in the 5-fold CV data set from the K562 cell line and the test data from the GM12878 cell line. For RFECS, the AUC values were not computed on the K562 CV data. The method with the best performance on the K562 or the GM12878 data were indicated with the bold font.

| Method | Cell line | AUC |
| --- | --- | --- |
| PREPRINT Bayesian | K562 | **0.993** |
| PREPRINT ML | K562 | 0.990 |
| PREPRINT Bayesian | GM12878 | **0.981** |
| PREPRINT ML | GM12878 | 0.978 |
| RFECS | GM12878 | 0.963 |

### PREPRINT predicted a larger number and shorter enhancers than RFECS and ChromHMM

PREPRINT and RFECS trained on the whole K562 training data predicted enhancers genome-wide in both cell lines, K562 and GM12878. Both PREPRINT and RFECS scanned the genome in subsequent 2 kb windows advancing in 100 bp shifts along the genome. For each of the genomic windows, PREPRINT and RFECS assigned a prediction score. If the prediction score exceeded a certain threshold, the window was predicted as an enhancer. Choosing the prediction threshold for a classifier is a critical task. Therefore, as a first choice, a prediction threshold of 0.5 was adopted by both PREPRINT and RFECS. For PREPRINT this implied that in order to predict a genomic region as an enhancer, the enhancer class probability needed to exceed 0.5. In contrast, in order for RFECS to predict a genomic region as an enhancer, 50% or more of the trees of the random forest needed to vote for an enhancer class. However, the prediction threshold of 0.5 was likely suboptimal and not well calibrated. Hence, for PREPRINT, the best operating point threshold and the 1% false positive rate (FPR) threshold were estimated from the performance evaluation measures on the K562 CV data set. In addition, the best operating point threshold and the 1% FPR threshold were estimated from the performance evaluation measures on the GM12878 test data. In both cases, the classifiers were trained on the K562 data. It must be stressed that the prediction thresholds estimated from the K562 training data should be adopted when predicting enhancers in other cell lines, since the test regions might be unavailable for the other cell lines. The best operating thresholds, the 1% FPR thresholds as well as the number of enhancers predicted by the different methods with varying thresholds are listed in Table 2. The number of predictions was conditional on the chosen method and the adopted threshold. In the following sections, the method comparison results are often presented for an equal number of enhancer predictions obtained by PREPRINT and RFECS. Therefore, to equalise the number of enhancers predicted by RFECS and PREPRINT for comparison purposes, the prediction thresholds of the methods were adjusted so that both methods predicted the same number of enhancers.

**Table 2:** The number of genome-wide enhancers predicted by different methods and thresholds.

| Method | Cell | The threshold of 0.5 |  | The best operating point or the threshold of 0.25 for RFECS |  |  | The FPR 1% Threshold |  |  |
| --- | --- | --- | --- | --- | --- | --- | --- | --- | --- |
|  |  | all | without TSS | threshold | all | without TSS | threshold | all | without TSS |
| PREPRINT ML | K562 | 27347 | 21238 | 0.51917 | 26423 | 20476 | 0.65472 | 20531 | 15531 |
| PREPRINT Bayesian | K562 | 36079 | 29747 | 0.35121 | 45472 | 37387 | 0.63137 | 29632 | 24553 |
| RFECS | K562 | 18956 | 15786 | 0.25 | 44522 | 35089 |  |  |  |
| ChromHMM Weak Enhancer | K562 | 180471 | 176912 |  |  |  |  |  |  |
| ChromHMM Strong Enhancer | K562 | 69019 | 66888 |  |  |  |  |  |  |
| PREPRINT ML | GM12878 | 34448 | 29189 | 0.51917 | 33358 | 28246 | 0.65472 | 26274 | 22088 |
| PREPRINT Bayesian | GM12878 | 42480 | 38048 | 0.35121 | 52176 | 46446 | 0.63137 | 35447 | 31904 |
| RFECS | GM12878 | 20059 | 17746 | 0.25 | 58560 | 49699 |  |  |  |
| ChromHMM Weak Enhancer | GM12878 | 178474 | 175487 |  |  |  |  |  |  |
| ChromHMM Strong Enhancer | GM12878 | 64052 | 62599 |  |  |  |  |  |  |
| The thresholds estimated from the GM12878 test data |  |  |  |  |  |  |  |  |  |
| PREPRINT ML | GM12878 |  |  | 0.23675 | 58053 | 50012 | 0.68115 | 24934 | 20916 |
| PREPRINT Bayesian | GM12878 |  |  | 0.26116 | 60412 | 53681 | 0.81953 | 25401 | 23134 |

Predicting enhancers with PREPRINT and RFECS resulted in subsequent windows with a prediction score higher than the chosen threshold. To increase the resolution of the predictions, one or several single windows needed to be chosen within a wider region. RFECS predicted very wide regions and aimed to find multiple local maxima within a region. In turn, PREPRINT chose only one window with the maximum prediction score, and if multiple windows received the same maximum score, one was selected at random. Instead of predicting an enhancer as a single window, the whole region of subsequent enhancer predictions could be considered as an enhancer. Consequently, the proportions of the prediction region lengths were computed for the different enhancer prediction methods. The RFECS and PREPRINT predictions were produced with two different thresholds, 0.5 and 0.75. In addition to PREPRINT and RFECS, ChromHMM Strong Enhancer and Weak Enhancer clusters obtained from ENCODE were included in the method comparison [25, 26]. The proportions of the prediction lengths in the K562 cell line are illustrated in Figure 1. Figure 1 indicates that PREPRINT and RFECS predicted proportionally shorter enhancers than ChromHMM, ChromHMM predicting mostly enhancers with lengths varying from 1 kb to 10 kb. RFECS predicted a high proportion of enhancers with a length of 100 bp. In contrast, PREPRINT predicted proportionally more enhancers with lengths varying from 200 bp to 1 kb. In addition, PREPRINT Bayesian and RFECS predicted proportionally more enhancers with lengths larger than 1 kb compared to PREPRINT ML. Hereafter, the enhancers with lengths 100–1000 bp are referred to as short enhancers, and the enhancers with a length larger than 1 kb are referred to as long enhancers. The proportions of short and long enhancers predicted by RFECS were increased and decreased, respectively, when adopting the more stringent threshold of 0.75. This behaviour was expected because with the more stringent threshold, large prediction regions were divided into separate smaller regions and/or became shortened. In contrast, when adopting the more stringent threshold for PREPRINT, the distribution of the length frequencies remained almost the same. This suggests that the prediction scores of PREPRINT advanced from a low value to a high value and back within a short region (within a low number of window shifts) whereas the prediction scores of RFECS increased and decreased smoothly within a large genomic window. To conclude, the lengths of the PREPRINT enhancers were less sensitive to changes in the prediction threshold compared to the RFECS enhancers. Similar results were obtained for the genome-wide enhancers predicted on the GM12878 data (Supplementary Figure S4, Additional File 1). In comparison to Figure 1, the differences between the methods in the normalised frequencies of short and long enhancers were even more evident in the GM12878 cell line. Finally, a large proportion of PREPRINT enhancers consisted of only one window (100 bp) or of short enhancers (*<* 2 kb). This suggests that choosing the window with the maximal score within a larger prediction region was an adequate approach to define the exact locations of the PREPRINT enhancers.

**Figure 1:**
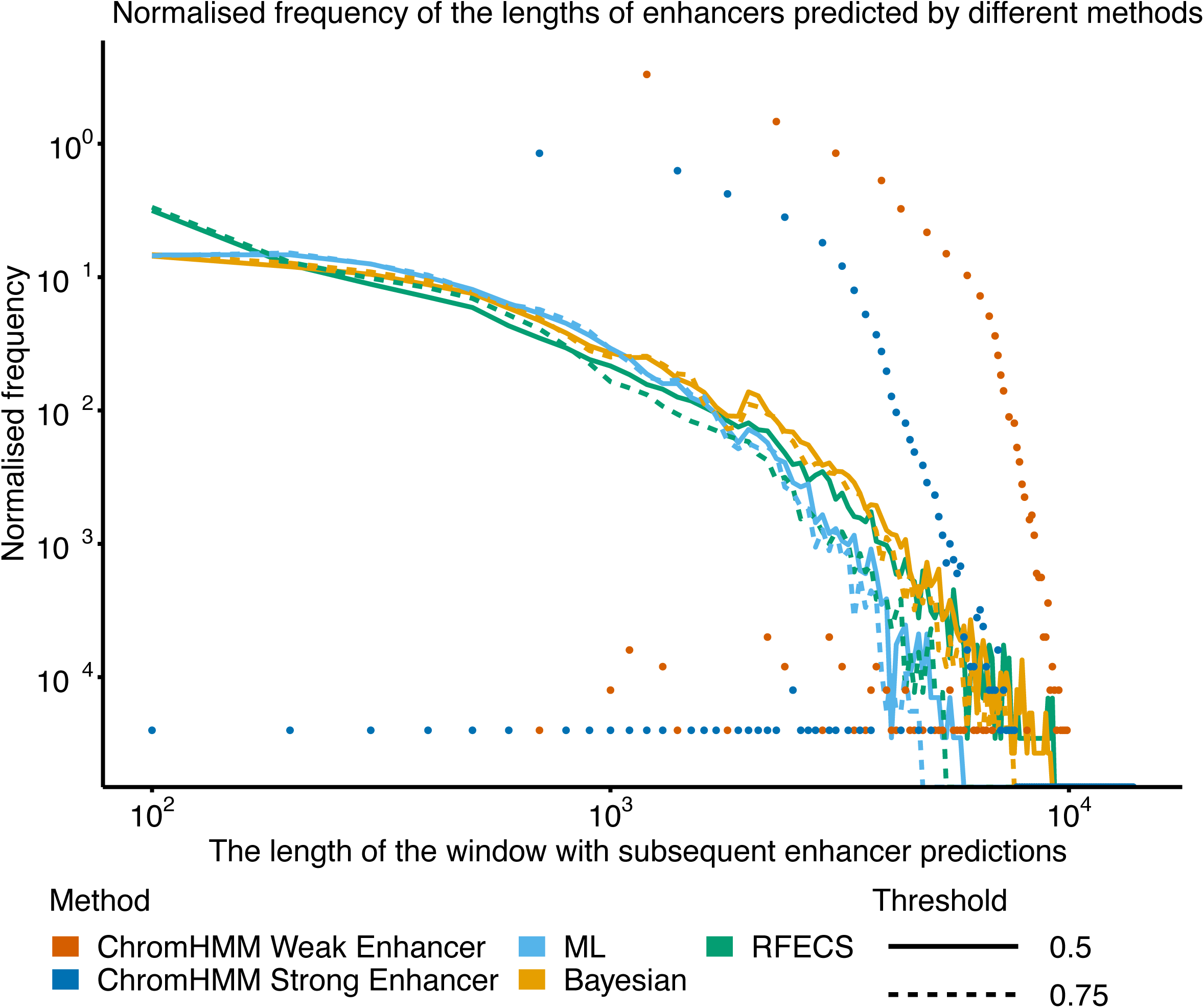
The normalised frequencies of the enhancer lengths. The enhancers were predicted in the K562 cell line by PREPRINT and RFECS with the thresholds of 0.5 and 0.75. For each method and threshold, the frequencies were divided by the total number of regions predicted as enhancers. The regions were formed by combining the subsequent enhancer predictions into a single region.

The number of genome-wide enhancer predictions obtained by PREPRINT, RFECS and ChromHMM in both cell lines K562 and GM12878 are provided in Table 2. The number of predictions were recorded before and after TSS removal. In addition, the number of predictions obtained by PREPRINT and RFECS with varying thresholds were reported. The RFECS predictions were obtained with the prediction thresholds of 0.5 and 0.25. The PREPRINT enhancers were obtained with the prediction threshold of 0.5, the best operating point threshold, and the 1 % FPR threshold. The numbers of PREPRINT predictions in the GM12878 cell line were obtained either utilising the thresholds estimated from the K562 CV data or the thresholds estimated from the GM12878 data. As a result, the best operating point threshold for PREPRINT ML in the K562 cell line were close to 0.5 whereas the best operating point thresholds for the PREPRINT Bayesian as well as the thresholds estimated on the GM1287 data ranged from 0.2 to 0.3. These lower best operating point thresholds resulted in significantly larger number of enhancers compared to the threshold of 0.5. Finally, the 1% FPR thresholds estimated from the GM12878 test data were more stringent compared to the 1% FPR thresholds estimated from the K562 CV data, especially for the PREPRINT Bayesian approach. The observed differences in the prediction thresholds and hence in the prediction numbers suggest a need for a careful calibration of the prediction thresholds. Especially, additional caution is required when generalising the prediction thresholds between data originating from different cell lines.

Overall, the number of enhancers predicted by RFECS with the prediction threshold of 0.5 was lower than the numbers predicted by PREPRINT with the threshold of 0.5 and by ChromHMM. The prediction numbers obtained by PREPRINT with the 1% FPR threshold became comparable with the prediction numbers obtained by RFECS with the prediction threshold of 0.5. Accordingly, the number of RFECS enhancers obtained with the lower prediction threshold of 0.25 became comparable with the number of enhancers obtained by PREPRINT with the threshold of 0.5. In addition, the Bayesian approach predicted a higher number of enhancers than the ML approach. The larger number of enhancer predictions for PREPRINT may result from PREPRINT predicting proportionally more short enhancers than RFECS, as seen, for example, in Figure 1 and Supplementary Figure S4, Additional File 1. The larger number of PREPRINT enhancers may also result from PREPRINT prediction score fluctuating between the enhancer and non-enhancer class unnecessarily often within a short stretch of DNA.

For comparison, the number of genome-wide enhancer predictions and thresholds for the different methods trained separately on the two random region definitions are provided in Supplementary Table S2 Additional File 1. The number of enhancer predictions were notably higher compared to the number of predictions obtained by the methods trained on the combined random regions. Especially, the methods trained on the pure random regions predicted a substantial number of enhancers, even with the 1% FPR thresholds. In contrast, the methods trained on the combined random regions resulted in an adequate classification performance and in a sensible number of genome-wide enhancer predictions. While studying further the differences between the results obtained by the methods on varying training data definitions would be interesting, the results in the following sections are presented only for the methods trained on the combined random regions.

### Proportions of predictions overlapping a varying number of transcription factor and co-regulatory ChIP-seq peaks

The genome-wide enhancer predictions were validated by first inspecting the overlap between the predicted enhancers and the histone acetyltransferase (p300) binding sites (SydhK562P300Iggrab and SydhGm12878P300Iggmus ChIP-seq peak sets produced by ENCODE). In addition, a large set of TF and other co-regulatory protein binding sites from the Transcription factor ChIP-seq Uniform peaks produced by ENCODE were utilised for validation. The peaks for RNA Pol II, CTCF, CREB-binding protein (CBP) and p300 were removed from the Uniform peak set resulting in the peak sets of 111 and 76 individual DNA binding proteins for cell lines K562 and GM12878, respectively. The TF and co-regulatory factors are referred to as transcription related factors (TRF). For more details on the validation data, see Additional File 2.

As mentioned in the Background section, the co-occurrence of multiple TRF ChIP-seq peaks could be strong evidence for the transcriptional regulatory potential of the predicted enhancer. Thus, the first step in the validation was to investigate the number of overlapping TRF binding sites at the genome-wide enhancer predictions. An enhancer prediction overlapped a TRF binding site if the 2 kb prediction window overlapped with at least 1 bp of the TRF ChIP-seq peak. For each enhancer prediction obtained by the different methods, the number of overlapping TRF ChIP-seq peaks were computed. When computing the number of TRF ChIP-seq peaks overlapping a given enhancer, the individual TRF ChIP-seq peaks were not required to overlap with each other. The proportions of enhancer predictions overlapping a varying number of TRF binding sites in the K562 cell line are illustrated in Figure 2. In both comparisons **a** and **b** in Figure 2), an equal number of enhancers were predicted by PREPRINT and RFECS. In comparison **a**, the number of enhancers were 35089, the number of enhancers predicted by RFECS with the threshold of 0.25. In comparison **b**, the number 15531 corresponded to the number of enhancers predicted by PREPRINT ML with the 1% FPR threshold. Thus, in comparison **a**, a larger number of top enhancer predictions were considered, and notably the proportion of predictions without any TRF binding site were smaller for PREPRINT Bayesian than for RFECS and PREPRINT ML. In addition, the proportions of predictions overlapping 1–5 TRF binding sites were slightly higher for the PREPRINT methods. For the larger number of TRF binding sites (*>* 5), all methods demonstrated comparable proportions of overlap. In contrast, in comparison **b**, a smaller set of enhancer predictions were considered. The RFECS achieved the smallest proportion of predictions lacking a TRF binding site as well as the largest proportions of predictions with TRF binding sites ranging from 10 to 20. Conversely, the proportions of enhancers overlapping a small number (1–6) of TRF binding sites were higher for PREPRINT, especially for the PREPRINT Bayesian approach.

**Figure 2:**
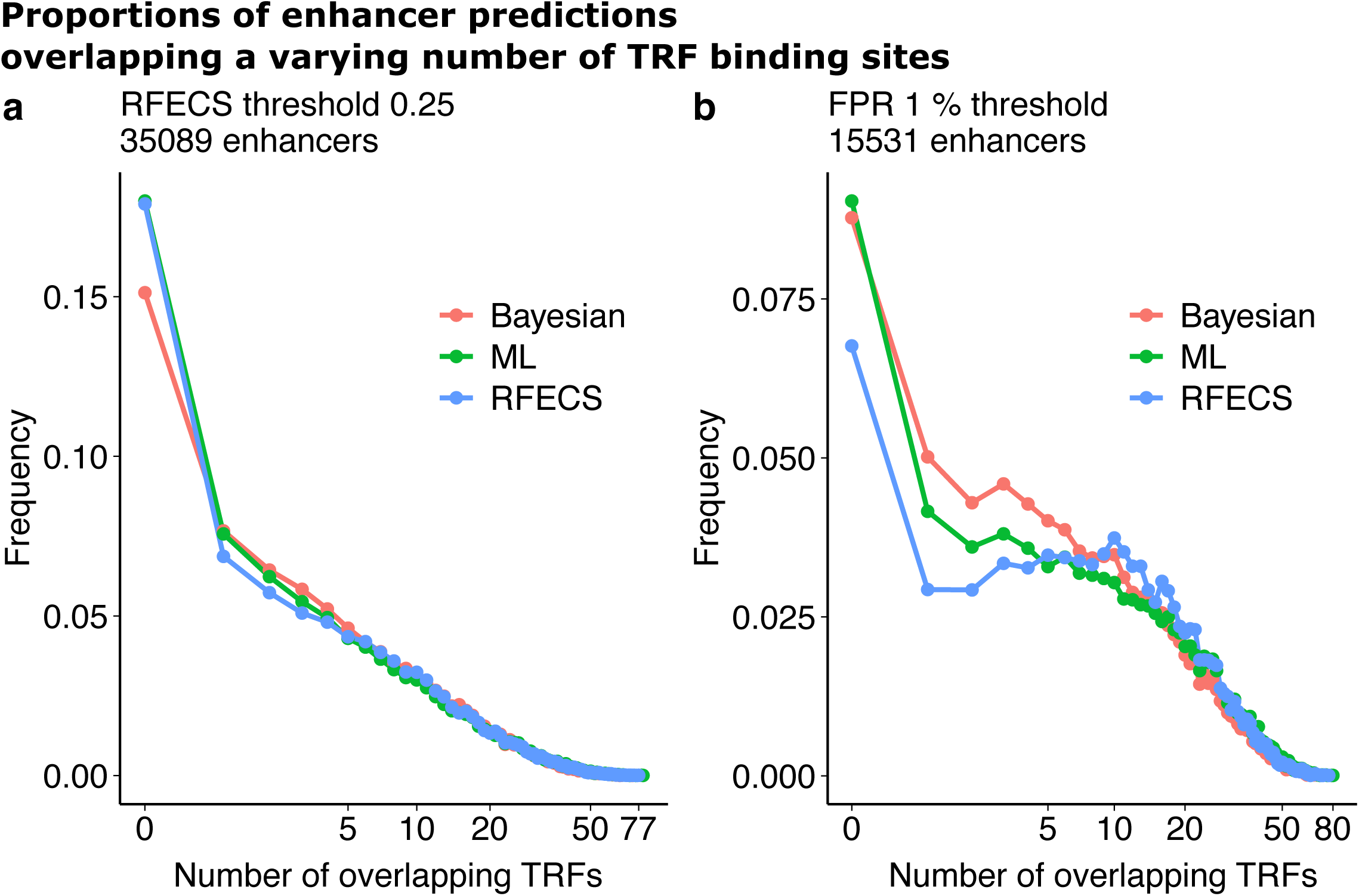
The proportions of the genome-wide enhancer predictions overlapping the varying number of TRF ChIP-seq peaks in the K562 cell line. The number of enhancers in each comparison are shown above the figure. In comparison **a**, the number of enhancers was the number of enhancers predicted by RFECS with the threshold of 0.25, and in comparison **b**, the number of enhancers was the minimum number of enhancers predicted by PREPRINT methods with their 1% FPR thresholds.

Similar proportions of enhancer predictions overlapping a varying number of TRF binding sites were obtained for the predictions in cell line GM12878 (see Supplementary Figure S5, Additional File 1). In comparison **a**, the number of enhancers was the number of predictions obtained by RFECS with the threshold of 0.25 (49699), and in comparison **b**, the number of enhancers was the number of predictions obtained by the PREPRINT ML approach with the 1% FPR threshold estimated on the K562 CV data (22088). Again, with the larger set of enhancers in Figure S5**a**, the proportion of predictions lacking a TRF binding site were smaller for the PREPRINT methods than for RFECS. Moreover, the proportions of predictions overlapping from 1 to 10 TRF binding sites were higher for PREPRINT than for RFECS. Considering a smaller set of enhancer predictions in Supplementary Figure S5**b**, Additional File 1, the proportion of predictions lacking a TRF binding site were higher for PREPRINT than for RFECS. Nevertheless, the proportions of enhancers overlapping a small number (1–3) of TRF binding sites were higher for PREPRINT than for RFECS, the PREPRINT Bayesian achieving higher proportions that PREPRINT ML.

To conclude, the proportions of PREPRINT predictions overlapping a varying number of TRF binding sites demonstrated desirable distributions. Firstly, the low proportion of predictions lacking a TRF binding site would be a preferable result for a set of enhancer predictions. According to the results, the proportion of predictions lacking a TRF binding site was lower for the PREPRINT methods than for RFECS when requiring a large set of predictions by relaxing the prediction threshold. Secondly, PREPRINT enhancers contained proportionally more enhancers overlapping 1–10 different TRF peaks than the predictions obtained by RFECS; these enhancers may have displayed weaker chromatin feature signals and may have been missed by RFECS, while they were still validated according to the TRF ChIP-seq data. Thirdly, the proportion distributions between RFECS and PREPRINT were comparable when requiring a large number (*>* 10) of TRFs overlapping the enhancer predictions. In addition, the proportions between methods became comparable when the number of predictions increased. Finally, as stated above, the co-occurrence of multiple TRF ChIP-seq peaks could indicate strong validation evidence of the predicted enhancer. However, defining the threshold for the number of overlapping TRF peaks is not straightforward, for example, according to the results presented here. Therefore, instead of requiring a more stringent thresholds for the number of overlapping TRF peaks, the most relaxed setting possible was adopted for the validation. In the results presented in the following sections, an enhancer prediction was validated if at least 1 bp of the 2 kb prediction window overlapped with at least 1 bp of at least 1 TRF ChIP-seq peak. Similar validation procedure was adopted to the 2 kb RFECS predictions and the ChromHMM predictions of varying lengths.

### The validation of genome-wide predictions

To study the performance of the methods to predict enhancers in the whole human genome, again an equal number of enhancers predicted by PREPRINT and RFECS were selected. In addition to the enhancer predictions, an equal number of non-enhancer predictions were chosen. The non-enhancers were sampled randomly from the regions which obtained a prediction score less or equal to 0.5. For both RFECS and PREPRINT, the number of enhancers and non-enhancers was set to the number of enhancers predicted by RFECS with the threshold of 0.25 or to the number predicted by PREPRINT methods with the 1 % FPR threshold. The number were the same numbers of enhancers as in the comparisons illustrated in Figure 2 and Supplementary Figure S5, Additional File 1. The predicted enhancers and non-enhancers were labelled either as true positives, false positives, true negatives or false negatives considering the overlap between the regions and the TRF ChIP-seq peaks. The TRF ChIP-seq peaks were divided into two validation sets: the p300 ChIP-seq peaks and the TRF ChIP-seq peaks containing all other peaks except the p300 peaks. The genome-wide enhancer and non-enhancer predictions with the the validation information were utilised to compute the AUC values. The AUC values are provided in Table 3. As a result, RFECS achieved the highest AUC scores in all settings. Nevertheless, PREPRINT achieved comparable AUC values to RFECS when considering the TRF validation data, especially in the GM12878 cell line. Of the PREPRINT methods, the Bayesian approach outperformed the ML approach in most of the comparisons (5 out of 8). To conclude, the validation performance of the genome-wide predictions was comparable across methods, and the PREPRINT achieved a good generalisation performance between data from different cell lines. However, the various settings, e.g. the cell line adopted for training and prediction, the type of validation data, and the number of enhancers and non-enhancers led to divergent results.

**Table 3:** The AUC values for the genome-wide predictions. The true labels of the predictions were based on the overlap between the predictions and the TRF ChIP-seq peaks. An equal number of enhancers predicted by PREPRINT and RFECS were chosen; the numbers are indicated in the column #enhancer predictions. In each combination of the validation data (p300 or TF) and the method, the AUC value for the best method was highlighted with the bold font. In addition, the AUC values for PREPRINT predictions in cell line GM12878, computed utilising the TRF validation data, were highlighted due to a comparable generalisation performance to RFECS.

| Cell line | method | #enhancer predictions | p300 | TRF |
| --- | --- | --- | --- | --- |
| K562 | PREPRINT ML | 35089 | 0.857 | 0.874 |
| K562 | PREPRINT Bayesian | 35089 | 0.855 | 0.887 |
| K562 | RFECS | 35089 | **0.911** | **0.900** |
| K562 | PREPRINT ML | 15531 | 0.850 | 0.898 |
| K562 | PREPRINT Bayesian | 15531 | 0.856 | 0.908 |
| K562 | RFECS | 15531 | **0.912** | **0.929** |
| GM12878 | PREPRINT ML | 49699 | 0.826 | **0.889** |
| GM12878 | PREPRINT Bayesian | 49699 | 0.833 | **0.888** |
| GM12878 | RFECS | 49699 | **0.911** | **0.895** |
| GM12878 | PREPRINT ML | 22088 | 0.821 | **0.911** |
| GM12878 | PREPRINT Bayesian | 22088 | 0.826 | **0.910** |
| GM12878 | RFECS | 22088 | **0.881** | **0.941** |

To further investigate the performance of the methods to predict the enhancers genome-wide, the proportion of validated enhancers, e.g., the validation rate, was computed for a varying number of the top genome-wide predictions. The validation rates for the predictions obtained in the K562 cell line by the different methods are illustrated in Figure 3. In Figure 3, the numbers on the x-axis correspond to the number of enhancers predicted by: RFECS with the prediction threshold of 0.5 (15786), PREPRINT ML with the threshold 0.5 of (21238), PREPRINT Bayesian with the threshold of 0.5 (29747), and RFECS with the threshold of 0.25 (35089). In addition to computing the validation rate for the given set of enhancer predictions, the validation rates were computed for a random region set of equal size (yellow bars in Figure 3). For comparison, the validation rates were also provided for the ChromHMM Weak and Strong Enhancers clusters. To conclude, the validation rates of enhancers predicted by any of the methods were clearly higher than the validation rates of the random regions. When comparing the different methods, the validation rates were higher for RFECS than for PREPRINT with a low number of enhancer predictions (15786 and 21238). Nevertheless, when considering a high number of enhancers (29747 and higher), PREPRINT achieved comparable or better validation rates than RFECS. Notably, for the high number of enhancers, the predictions obtained by PREPRINT Bayesian achieved a higher validation rate than the predictions obtained by RFECS.

**Figure 3:**
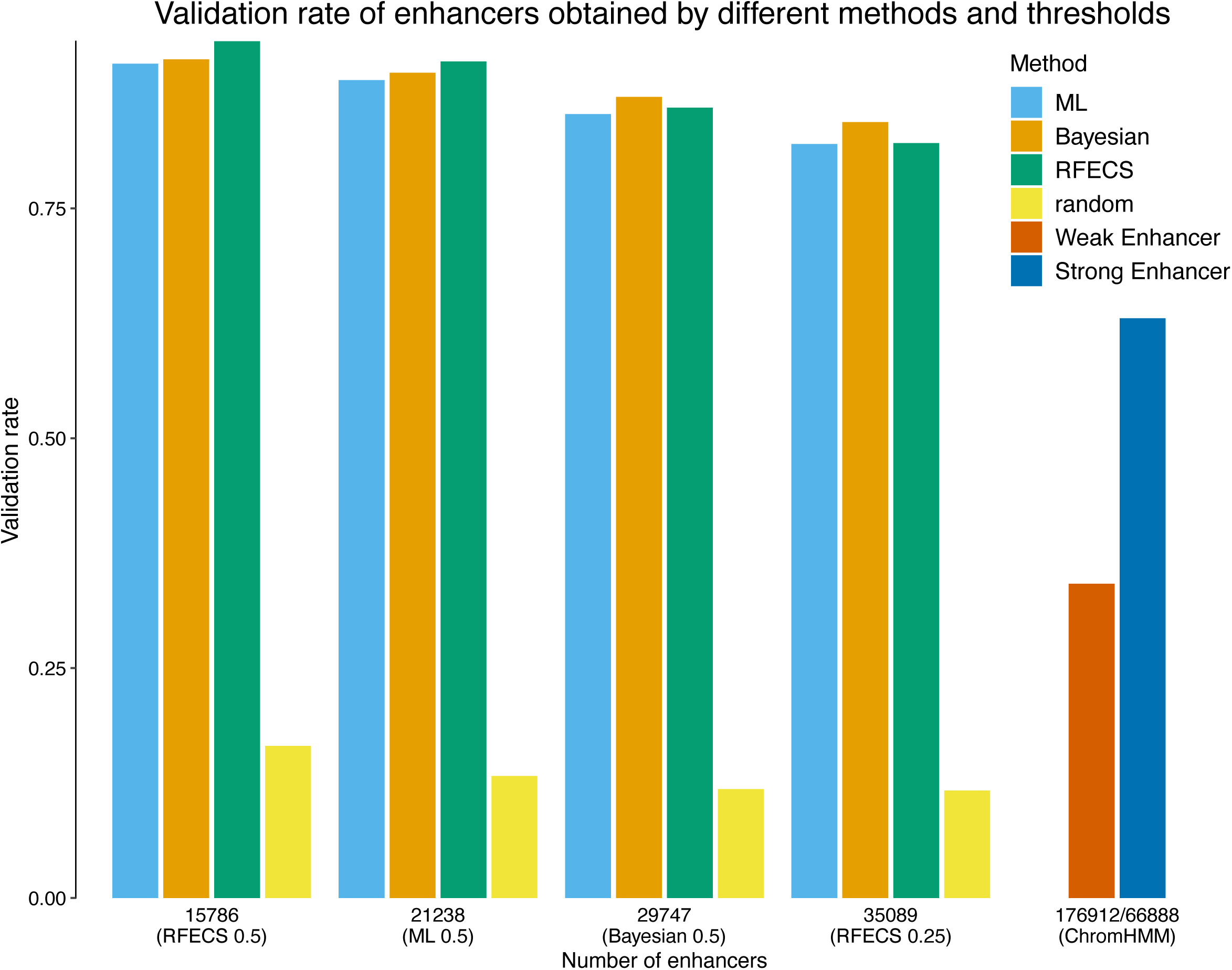
The validation rate of the genome-wide enhancer predictions obtained by the different methods and thresholds in the cell line K562. An enhancer prediction was validated if the 2 kb prediction window overlapped with at least 1 bp of at least one TRF ChIP-seq peak.

The validation rates for the predictions obtained in the GM12878 cell line by the different methods are demonstrated in Supplementary Figure S6, Additional File 1. Considering the smallest set of top predictions (17746), RFECS achieved the highest validation rate. However, with the larger set of predictions (38048 and larger), the validation rate of RFECS dropped rapidly whereas the PREPRINT methods stayed at the same validation rate level. In the GM12878 cell line, the difference between the performance of the PREPRINT methods was negligible. These results suggest that RFECS may have predicted well a restricted set of enhancers with the strongest chromatin feature signals whereas PREPRINT predicted a larger number of enhancers, containing enhancers with both strong and weak feature signals. When requiring the methods to predict a large number of enhancers, the PREPRINT enhancers had a higher validation rate compared to RFECS and ChromHMM enhancers. Moreover, PREPRINT trained on the K562 data generalised well on data from the GM12878 cell line.

### The overlap between enhancers predicted by different methods

The overlaps between the enhancers predicted by the different methods were investigated. The overlap was defined as follows: PREPRINT and RFECS predictions were again considered as 2 kb genomic regions whereas the ChromHMM enhancers were regions of varying lengths. For any two enhancers predicted by any two separate methods to overlap, at least 1 bp overlap was required. As an enhancer predicted by the one method might overlap with two enhancers predicted by the other method, the overlap between enhancers predicted by a pair of methods were not symmetrical. Hence, the overlap was computed in both directions. The numbers of overlapping genome-wide predictions obtained by the different methods as well as the numbers of the unique enhancer predictions for each method were illustrated as Venn diagrams. In each region or intersection set defined by the Venn circle curves, the number of enhancers belonging to that set was provided together with the percentage of validated enhancers in that set. The validation was again performed as described above.

The Venn diagram of the predictions obtained in the K562 cell line is illustrated in Figure 4. In comparisons **a** and **c**, the PREPRINT predictions were obtained by the ML approach whereas in **b** and **d**, the predictions were obtained by the Bayesian approach. In each comparison, the number of PREPRINT and RFECS enhancers was equal. In comparisons **a** and **b**, the numbers were chosen as the minimum number of enhancers predicted by RFECS with the threshold of 0.5 and PREPRINT methods with the 1% FPR threshold. The numbers were 15531 and 15787, respectively. In comparisons **c** and **d**, the numbers were the number of enhancers predicted by RFECS with the prediction threshold of 0.25 (35089). Overall, around half of the enhancers predicted by PREPRINT or RFECS were predicted by all of the three methods, and this intersection set had the highest validation rate (90% or larger). In addition, in the intersection set, the number of enhancers predicted by ChromHMM were much larger than for PREPRINT or RFECS. Hence, the validation rate of ChromHMM enhancers in the intersection set was lower (around 70%), likely reflecting the fact that ChromHMM enhancers were imprecise and/or the cluster labels along the genome alternated frequently between the enhancer state and the other states. Furthermore, the enhancer predictions shared by any two methods attained high validation rates (60–90%), and for the enhancers predicted uniquely by a single method, the validation rate ranged from 50 to 90%. Of the enhancers uniquely predicted by the different methods, the RFECS enhancers tended to achieve the highest validation rate. Nevertheless, the unique enhancers predicted by PREPRINT Bayesian reached a comparable validation rate (64%) to the unique enhancers predicted by RFECS (65%) when considering the larger set of enhancer predictions (Figure 4**d**).

**Figure 4:**
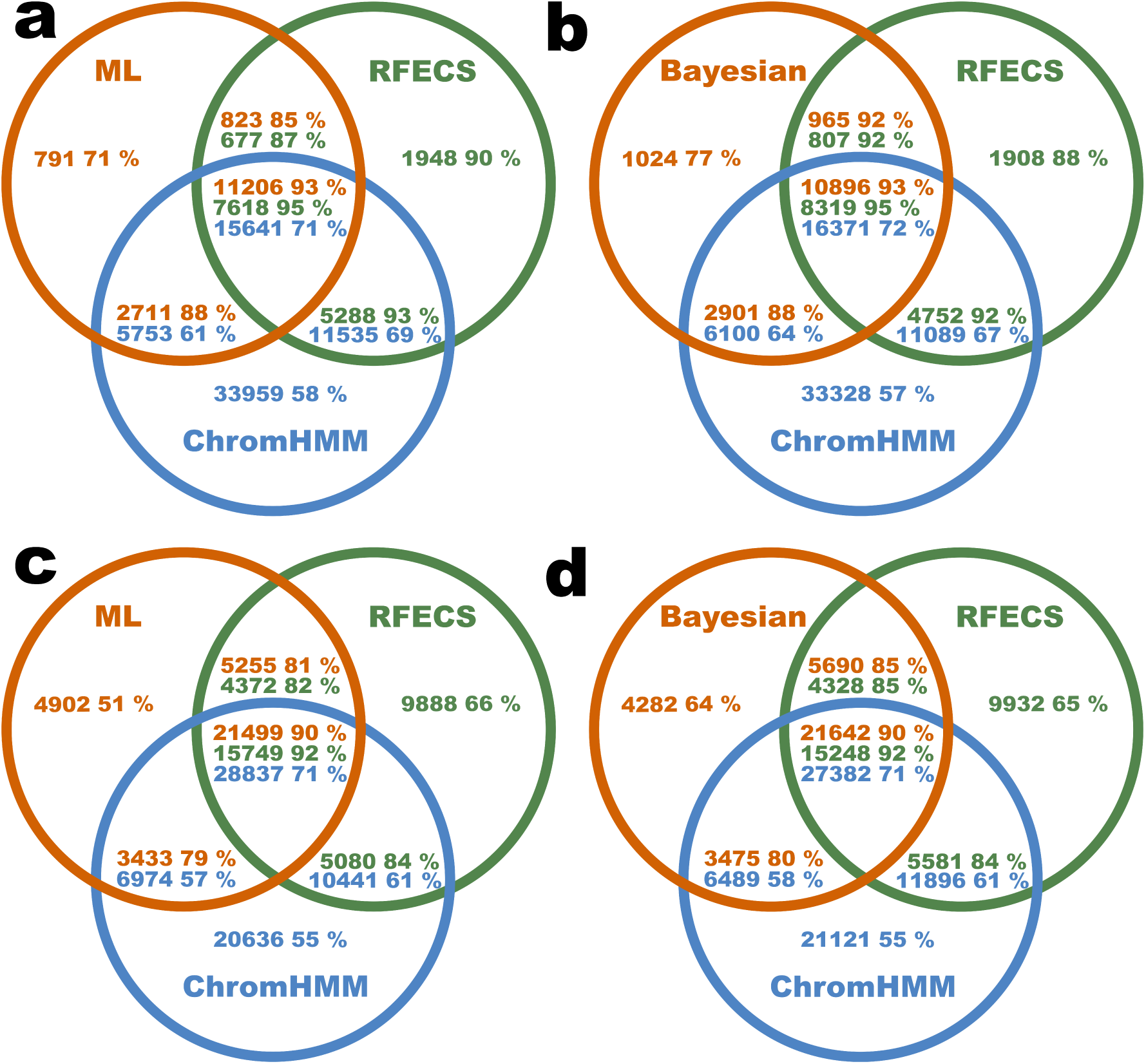
The unique and overlapping genome-wide enhancer predictions obtained by different methods in cell line K562. In comparisons **a** and **c**, the PREPRINT predictions were obtained by the ML approach whereas in comparisons **b** and **d**, the PREPRINT predictions were obtained by the Bayesian approach. The overlap between the PREPRINT, RFECS and ChromHMM predictions were quantified as the number of enhancers. In each comparison, the number of enhancers predicted by PREPRINT and RFECS was equal. The numbers were: **a** 15531, **b** 15786, **c** 35089, and **d** 35089. Inside every region or intersection, the number of enhancers in the given the set is indicated together with the percentage of validated enhancers in the set. The areas of the intersection sets are not proportional to the number of overlapping regions due to the asymmetry of overlaps.

Considering the intersecting enhancers predicted by all of the three methods, PREPRINT predicted a larger number of enhancers than RFECS, implying that one RFECS prediction overlapped with multiple neighbouring PREPRINT predictions. Thus, PREPRINT behaved similarly as ChromHMM by predicting multiple enhancers along a short stretch of DNA. With relaxing the threshold (Figure 4**c** and **d**), the number of ChromHMM enhancers in the intersection set increased; this was a consequence of PREPRINT and RFECS beginning to cover more ChromHMM enhancers. Finally, PREPRINT methods predicted the smallest number of unique predictions in all comparisons, which could be seen as an advantage.

The Venn diagrams for the predictions in the GM12878 cell line are provided in Supplementary Figure S7, Additional File 1. In comparisons **a** and **c**, the PREPRINT predictions were obtained by the ML approach whereas in comparisons **b** and **d**, the predictions were obtained by the Bayesian approach. In each comparison, the number of PREPRINT and RFECS enhancers was equal. In comparisons **a** and **b**, the numbers were chosen as the minimum number of enhancers predicted by RFECS with the threshold of 0.5 and PREPRINT methods with the 1% FPR threshold. The numbers were both 17746. In comparisons **c** and **d**, the numbers were the number of enhancers predicted by RFECS with the prediction threshold of 0.25 (49699). The results in the GM12878 cell line were comparable to ones obtained in the K562 cell line. However, in all comparisons, the number of unique enhancers predicted by RFECS was considerably high, and their validation rate, especially when relaxing the prediction threshold (Supplementary Figure S7**c** and **d**), was low (30%). In contrast, when relaxing the prediction threshold, the validation rate of the unique PREPRINT predictions remained almost the same (50–60%).

Finally, to compare the overlap between predictions made by PREPRINT ML and PREPRINT Bayesian together with their overlap with RFECS predictions, the number of unique and intersecting predictions were computed. The resulting Venn diagrams are shown in Supplementary Figure S8, Additional File 1. The comparisons **a** and **b** illustrate the unique and intersecting genome-wide enhancers predicted by PREPRINT and RFECS in the K562 cell line whereas in comparisons **c** and **d**, the predictions were obtained in the GM12878 cell line. In each comparison, an equal number of top enhancer predictions was selected for each method, and the numbers were: **a** the minimum number of enhancers predicted by PREPRINT with 1% FPR threshold in the K562 cell line, **b** the number of enhancers predicted by RFECS with the threshold of 0.25 in the K562 cell line, **c** the minimum number of enhancers predicted by PREPRINT with the 1% FPR threshold in cell line GM12878, and **d** the number of enhancers predicted by RFECS with the threshold of 0.25 in the GM12878 cell line. Overall, again around half of the enhancers predicted by any of the method were found by all methods, and this intersection set achieved the highest validation rate (85–95%). In addition, there were significant numbers of enhancers predicted by any pair of methods. Notably, the validation rate of intersecting enhancers between PREPRINT and RFECS exceeded the validation rate of intersecting enhancers between the PREPRINT ML and Bayesian approach. Lastly, there were also significant numbers of enhancers predicted by one method only. RFECS predicted the highest number of unique enhancers achieving a high validation rate (88–91%) when considering a smaller number of enhancer predictions (Supplementary Figure S8**a** and **c**, Additional File 1). However, the validation rates of unique RFECS predictions were low when requiring a larger set of enhancer predictions, especially in the GM12878 cell line (37%). Of the unique enhancers predicted by PREPRINT, the predictions obtained by the Bayesian approach achieved the highest validation rate (70–85%).

As a conclusion, PREPRINT trained on the K562 data generalised successfully on the GM12878 data, and the Bayesian approach performed sufficiently. In some comparisons, the Bayesian approach attained similar or even superior performance to RFECS. When relaxing the prediction threshold, RFECS began to predict more enhancers not covered by the other methods, and these enhancers obtained a low validation rate. In contrast, PREPRINT predicted a low number of unique enhancers even when relaxing the prediction threshold; this may be a desirable property of PREPRINT. However, it was challenging to compare the validation performance of the overlaps between different methods due to various reason. The first reasons was the number of the unique enhancers varying greatly between the methods. Secondly, multiple PREPRINT enhancers tended to overlap with one RFECS enhancer, complicating the comparisons further. The asymmetric overlap was likely due to PREPRINT prediction scores fluctuating along the genome instead of changing smoothly as the RFECS prediction scores. Thirdly, RFECS required a distance of at least 2 kb between two subsequent enhancers. In contrast, PREPRINT did not involve such a requirement for the distance between predicted enhancers. Thus, PREPRINT tended to predict multiple individual enhancers within a short stretch of DNA. Predicting multiple individual enhancer within a short stretch of DNA also permitted two PREPRINT predictions to overlap with the same validating TRF ChIP-seq peak. This may have resulted in high validation rates for the PREPRINT predictions.

### Examples of validated enhancers uniquely predicted by PREPRINT

Some examples of validating enhancers uniquely predicted by PREPRINT were visualised in a genome browser together with the chromatin feature data. The examples of the predicted enhancers in a 20 kb genomic locus in chromosome 1 are demonstrated in Figure 5. The enhancers were predicted in the K562 cell line. Starting from the top, the validating predictions are indicated as red arrows. Below the arrows, the PREPRINT and RFECS predictions are provided, together with the predictions scores and the 0.5 threshold line. In addition, the ChromHMM predictions, the GENCODE genes, a subset of the chromatin feature ChIP-seq data tracks, and a subset of the validating Uniform TRF ChIP-seq peaks are displayed in Figure 5. The data tracks for all 15 chromatin features as well as the whole collection of the TRF Chip-seq peaks in the 20 kb locus are provided in Additional File 3.

**Figure 5:**
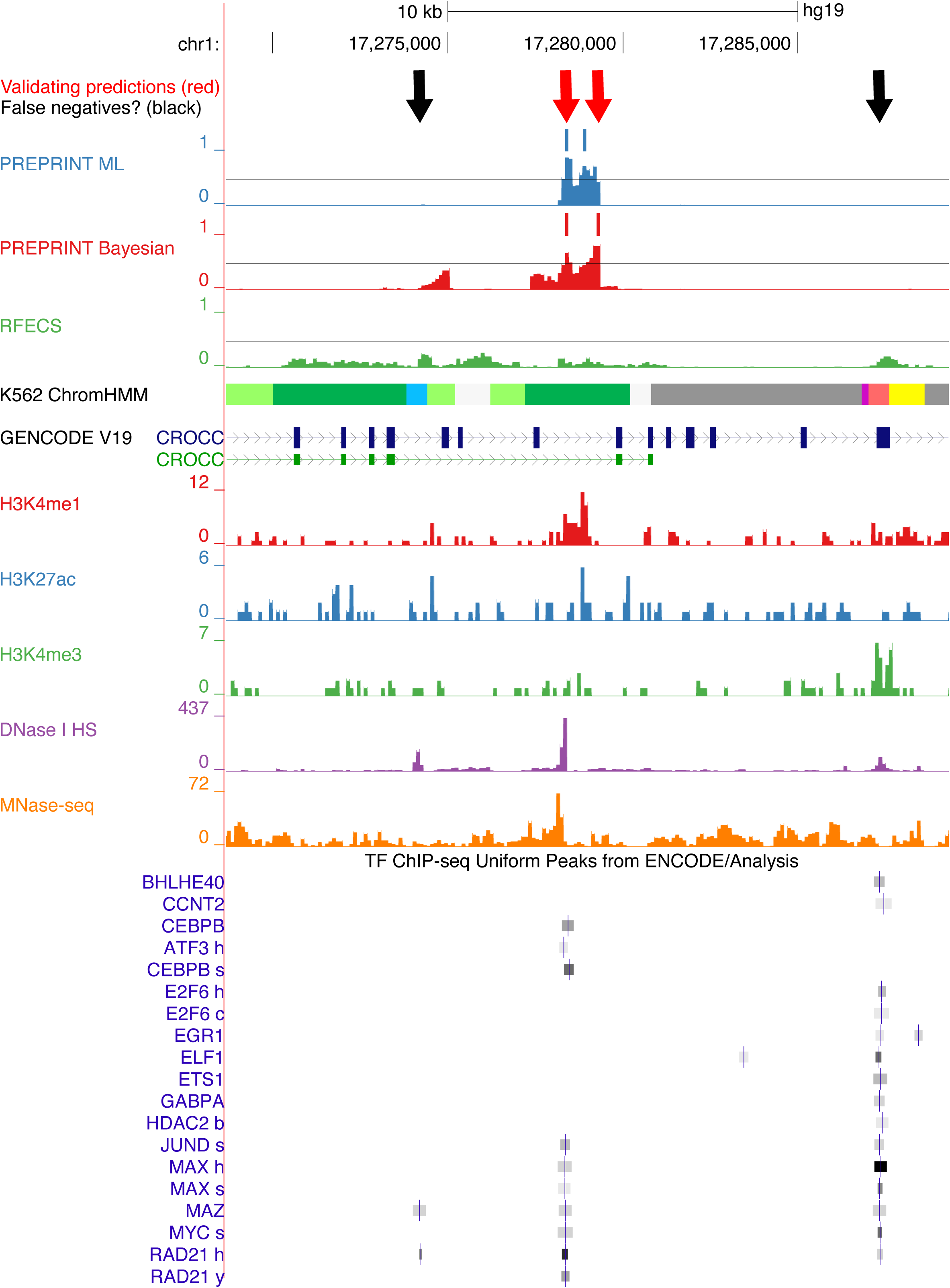
Genome browser visualisation for examples of enhancers uniquely predicted by PREPRINT. The data originated from cell line K562. Colour codes for ChromHMM clusters: light green: Weak transcription, dark green: Transcription elongation/transition, blue: Insulator, grey: Repetitive/Copy Number Variation (CNV) or repressed, purple: Poised promoter, light red: Weak promoter, yellow: Weak enhancer.

In the 20 kb locus illustrated in Figure 5, PREPRINT methods predicted two individual enhancers in an intron of a gene CROCC whereas RFECS predicts none. However, there is likely only 1 enhancer in the region indicated by the red arrows, according to the chromatin feature patterns and the localisation of the validating TRF ChIP-seq peaks. Moreover, the prediction scores of PREPRINT changed gradually from low values to high values in a stepped fashion, resulting in multiple predictions within a short stretch of DNA. Therefore, by lowering the prediction threshold, PREPRINT would predict only one enhancer at the location of the red arrows. Predicting multiple enhancers within a short stretch of DNA likely contributed to PREPRINT predicting a larger number of enhancers compared to RFECS with the threshold of 0.5. In addition, this behaviour likely resulted in a large number of PREPRINT predictions in the common set of enhancers predicted by all three methods, for example, as demonstrated in the Venn diagrams in Figure 4. Furthermore, predicting multiple adjacent enhancers caused the individual enhancers to overlap the same set of validating TRF ChIP-seq peaks, possibly producing overly high validation rates for the PREPRINT predictions. To further define the PREPRINT predictions, the two neighbouring predictions could be combined, for example, considering only one of them or their middle position.

In the 20 kb locus illustrated in Figure 5, there were also two possible enhancers indicated as black arrows. According to the TRF ChIP-seq peaks, they would validate as enhancers, hence they could be false negatives. At the leftmost black arrow, PREPRINT Bayesian reached prediction score just below the 0.5 threshold line, and the region at the rightmost black arrow was labelled as Weak promoter and Weak Enhancer by ChromHMM. All locations indicated by arrows have acquired DNase-seq peaks corresponding to open chromatin. To assure the enhancer function of these locations, one would need to perform in vivo functional enhancer assay.

Figure 6 demonstrates examples of enhancers predicted in the GM12878 cell line in a 40 kb intergenic genomic region in chromosome 4. The full set of data is provided in Additional File 4. In this genomic region, PREPRINT predicted uniquely two neighbouring enhancers indicated as red arrows. There indeed might be two individual enhancers indicated by the chromatin feature signals and the validating TRF ChIP-seq peaks. However, the PREPRINT prediction (the leftmost red arrow) was slightly off the exact location of the validating TRF ChIP-seq peaks (black arrow). The locus denoted as the black arrow displayed weak signals of H3K4me1, H3K27ac and DNase I HS, although was not clearly predicted as an enhancer by any of the methods. The RFECS did not predict any enhancers in this genomic window; the prediction scores for RFECS remained just below the 0.5 threshold line. ChromHMM labelled the region as heterochromatin.

**Figure 6:**
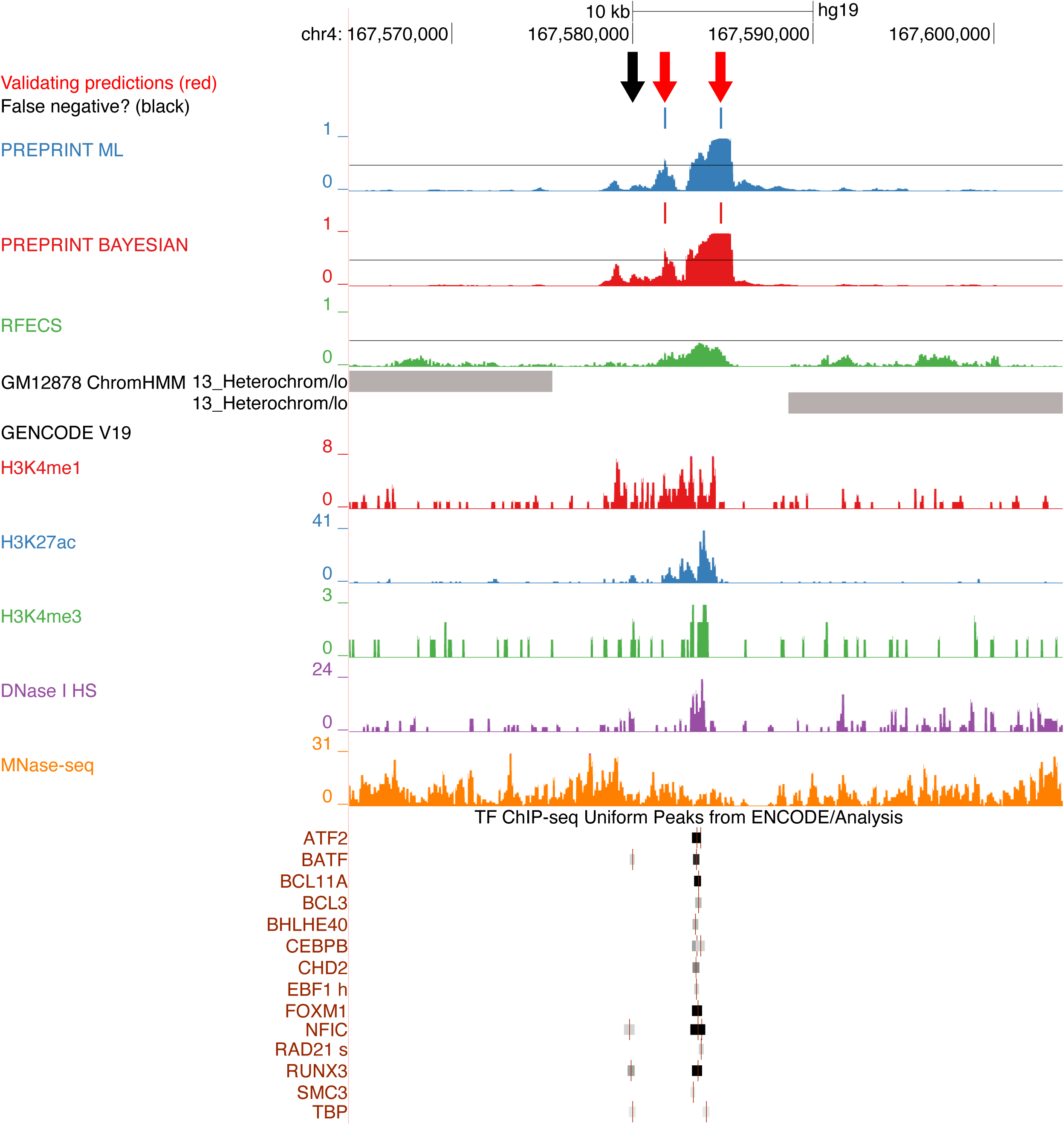
Genome browser visualisation for examples of enhancers uniquely predicted by PREPRINT. Data originated from cell line GM12878. Colour codes for ChromHMM clusters: Grey: Heterochromatin.

To conclude, PREPRINT predicted apparent enhancers not predicted by RFECS or ChromHMM. However, the visual inspection of the properties of the predictions reflected the results and challenges reported in the previous sections. One challenge is to predict enhancer locations with high accuracy. On one hand, the challenges are related to the definition of an enhancers and its boundaries. On the other hand, the challenges concern the choices adopted in the data analysis steps, such as the resolution of the data, the choice for the prediction threshold, the choice of the number of overlapping TRF ChIP-seq peaks for validation, and the definition of an overlap between two genomic regions or the set of regions. In other words, visual inspection of the individual predictions further justifies the concerns of being cautious when drawing conclusions about the enhancers predicted by machine learning methods.

## Discussion

This paper introduced PRobabilistic Enhancer PRedictIoN Tool (PREPRINT), a supervised enhancer prediction method based on probabilistic modelling of the characteristic chromatin feature patterns observed at enhancers and other genomic regions. PREPRINT recognises the dependency structure of the chromatin feature pattern associated with enhancers. In addition, PREPRINT models statistically the variability of the coverage patterns among individual enhancers. The statistical modelling provides robustness to the proposed approach and improves the generalisation ability of the classifier, for example, on data originating from cell line not utilised for training.

PREPRINT introduces two new probabilistic distance measures to quantify “the closeness” or “fit” of the genomic query regions and the characteristic patterns. Firstly, PREPRINT assumes Poisson distributed coverage counts with a scaled mean parameter. The scaled mean consists of the characteristic coverage pattern and a scaling parameter. Although an approximation, introducing the characteristic coverage pattern as an estimate of the Poisson mean incorporates the dependency structure of the feature patterns into the model. In addition, the scaling parameter accounts for the coverage signal variability due to various sources. The scaling parameters are estimated utilising two approaches: either sample-specific scaling parameters are estimated with a maximum likelihood (ML) approach or a global Gamma distribution is estimated for the scaling parameter (Bayesian approach). Furthermore, two probabilistic distance measures are defined depending on the approach employed to learn the scaling parameters. Hence, the two approaches are referred to as PREPRINT ML and PREPRINT Bayesian.

PREPRINT was trained and tested on the next-generation sequencing data produced by ENCODE for the myelogenous leukaemia cell line (K562) and the lymphoblastoid cell line (GM12878). Moreover, the prediction performance of PREPRINT was computationally compared to the state-of-the-art methods RFECS and ChromHMM. According to the results, the PREPRINT Bayesian trained on the data originating from the cell line K562 achieved the best performance on the test data originating from the GM12878 cell line.

To decrease the probability of misclassification for the whole-genome predictions, i.e., to control for the false positive rate, the prediction thresholds were calibrated. Adopting cross-validation within the training data, the threshold resulting in the 1% false positive rate (FPR) were obtained for PREPRINT. The threshold estimated from the K562 training data was assumed to generalise to the whole genome data and the GM12878 data. Therefore, the estimated threshold was utilized by PREPRINT to predict genome-wide enhancers in both cell lines. In addition, the threshold was altered to study the effect on the genome-wide predictions. Generally, the methods compared in this work predicted wide regions as enhancers. To pinpoint the individual enhancers within a wider prediction region, the methods compared in this study encountered challenges. Firstly, the prediction score threshold affected the width of the prediction region. For example, increasing the prediction threshold, one wide enhancer prediction region could be partitioned into two regions. Secondly, although RFECS predicted proportionally more short enhancers of length 100 bp compared to PREPRINT, the prediction scores of RFECS usually increased and decreased smoothly along the genome, resulting in very wide prediction windows and possibly uncertain predictions. Nevertheless, the prediction scores of PREPRINT advanced from a low value to a high value and back within a short region (within a low number of window shifts). As a result, the lengths of the PREPRINT enhancers were less sensitive to changes in the prediction threshold. However, PREPRINT was demonstrated to predict multiple enhancers within a short stretch of DNA.

The genome-wide enhancers predicted by the computational methods were validated with the largest collection to date of the TRF binding sites produced by the ENCODE consortia [43]. Firstly, the proportions of genome-wide predictions with a varying number of overlapping TRF ChIP-seq peaks were investigated. The proportions of predictions lacking a TRF binding site were lower for PREPRINT when requiring a larger set of prediction obtained by relaxing the prediction threshold. In addition, PREPRINT enhancers contained proportionally more enhancers overlapping 1–10 different TRF peaks than the predictions obtained by RFECS; these enhancers may have displayed weaker chromatin feature signals and been missed by RFECS, while they were still validated. Secondly, the validation rates of genome-wide enhancers obtained by different methods and prediction thresholds were compared. In this comparison, PREPRINT reached comparable or better performance to RFECS, especially when requiring the methods to predict a larger set of enhancers, i.e., employing a relaxed threshold. Moreover, the PREPRINT methods achieved a good generalisation performance between data from different cell lines. Finally, the common and unique set of enhancers predictions obtained by different methods and their validation rates were investigated. The unique enhancers predicted by PREPRINT Bayesian in cell line K562 reached a comparable validation rate (64%) to the unique enhancers predicted by RFECS (65%). The number of unique enhancers predicted by RFECS was considerably high, especially in cell line GM12878, and their validation rate, especially when relaxing the prediction threshold (Supplementary Figure S7**c** and **d**), was low (30%). When relaxing the prediction threshold, RFECS likely began to predict more enhancers not covered by the other methods, and these enhancers obtained a lower validation rate. In contrast, when relaxing the prediction threshold, the validation rate of the unique PREPRINT predictions remained almost the same (50–60%). However, PREPRINT behaved similarly as ChromHMM by predicting multiple enhancers along a short stretch of DNA. To conclude, PREPRINT trained on the K562 data generalised successfully on the GM12878 data, and the Bayesian approach performed sufficiently. In some comparisons, the Bayesian approach attained similar or even superior performance to RFECS. According to the results, PREPRINT Bayesian often outperformed the PREPRINT ML approach. This implies that using a global genome-wide distribution of the scaling parameter should be used, instead of modelling locally the fit of an individual sample, for example, to the enhancer aggregate pattern.

As stated above, the methods compared in this work predicted very wide genomic regions as enhancers. For a set of enhancer predictions to benefit, for example, the interpretation of SNPs, the exact enhancer location inside the wide prediction region need to be identified. In general, the concepts such as the exact enhancer location, length of an enhancer, the enhancer boundaries and the enhancer middle position are vague and need to be refined. An approach would be to define an enhancer as a stretch of DNA bounded by TRFs with the boundaries limited by the two well-positioned nucleosomes. To pinpoint the individual enhancers within a wider prediction region proved to be challenging. For example, PREPRINT often predicted two neighbouring enhancers (as in examples demonstrated in Figures 5 and 6). It is unclear whether these should be considered as separate enhancers, or whether any two neighbouring predictions should be combined. For example, the two enhancers could be merged by considering their middle location as the exact location of the enhancer. Finally, the resolution of the data, 100 bp adopted in this work, likely affected the widths of the predicted regions. Thus, increasing the data resolution could improve the resolution of the prediction.

For an enhancer to be validated, at least 1 bp overlap was required between an enhancer prediction window of size 2 kb and at least 1 TRF ChIP-seq peak. This definition of validation is not without problems: Firstly, a more rigorous requirement for the number of overlapping peaks could be adopted. However, to choose the threshold for the required number of overlapping peaks might not be straightforward and in this work, the most relaxed setting was adopted. In addition, instead of the prediction window of size 2 kb, a narrower prediction window, such as of 500 bp centred at the prediction location could have been utilized. Moreover, if two predictions were very close to each other, and both overlapped the same single TRF ChIP-seq peak, both were validated. Nevertheless, choosing the most relaxed setting is an adequate approach as the widths of the ChIP-seq peaks and the uncertainty associated to identifying the exact binding site of a TRF varies depending on many factors, such as the antibody specificity in the ChIP experiment, the TRF ChIP-seq data quality filtering, preprocessing of the raw reads, and the peak-calling methods. Some TRF ChIP-seq peaks might have been missed as they reside just below the selected significance threshold, often selected arbitrarily. Secondly, not all TSS-distal TRF binding sites are enhancers; they might be some other type of regulatory regions, such as insulators or silencers, or repressed enhancers. However, our training data does not support the identification of the latter. In addition, the TRFs may contain also repressive factors, not only activating. Finally, the human genome is estimated to encode around 1700 transcription factors [44] and a large number of co-regulatory factors whose binding sites in more than 200 different cell lines are largely unknown, for example, due to lack of antibodies. The co-occurrence of multiple TRF ChIP-seq peaks could be strong evidence for the transcriptional regulatory potential of the predicted region. However, this view is challenged by the recent work suggesting that these regions might be just technical artefacts of ChIP-seq, and the observation that these regions lack clear motifs [45, 46].

PREPRINT adopted a distribution resembling the Gamma-Poisson compound distribution to model the non-negative count data. However, different distributional assumptions could be considered. The exact negative binomial distribution [33, 34] could be utilised, or the count data could be corrected by the log-concave Poisson approach [36]. As the subtraction of the normalised control signal from the ChIP-seq signal resulted in continuous and negative values, instead of converting the data into non-negative counts and adopting the Gamma-Poisson compound model, Skellam [47] distribution or a distribution that models the difference between two independent negative binomial random variables [48] could have been employed. Moreover, the enhancer and non-enhancer aggregate patterns and the uncertainty inhered in them could be modelled, in contrast to assuming a fixed mean. However, this would lead to a more challenging estimating procedure, such as the expectation-maximisation (EM) algorithm. In contrast, RFECS does not involve any distributional assumptions and adapts readily to the continuous and real-valued ChIP-seq data. This can be an advantage for RFECS, as the data discretisation and especially the conversion of negative values to nonnegative both lose information. RFECS had access to the sharp negative dip of the chromatin feature coverage signals between the high positive signals from the two well-positioned nucleosomes, thus facilitating the separation of enhancers from the background coverage signal.

As demonstrated in this work, the choice for the random regions in the training data affected the performance of the methods. In addition to the random regions and promoters, other definitions of non-enhancers could be adopted, such as promoters driving different levels of gene expression, inactive promoters, exons, introns, miRNA and other non-coding genomic sites exhibiting at least some epigenetic signals. In addition, the feature patterns at individual non-enhancer regions could be scrambled to generate a versatile set of non-enhancer examples. Moreover, although not studied in this work, the choice of training data enhancer definition likely affects the results. To study the effect of the training data enhancer definition, the enhancers could be clustered to identify subclasses among them. The enhancers belonging to separate clusters might display different patterns and intensities of chromatin features [49, 50, 51]. It would be interesting to correlate the obtained clusters to biological and functional properties of the enhancers to identify enhancer subclasses or various enhancer states, such as poised, silenced, primed and active enhancers. The clustering could be done prior to the classifier training. Furthermore, in this work, the middle base of an enhancer was defined as the summit of the p300 ChIP-seq peak; this approach has naturally an inhered uncertainty associated with it. The individual ChIP-seq profiles could be shifted to improve the alignment between the enhancers. Moreover, the distribution of nucleosomes might vary between different enhancers, for example, the distance between the two well-positioned nucleosomes varies, and this should be considered in the prediction task.

With the advancements in the new types of sequencing data generation, big data analysis methods and machine learning, the race towards the ultimate genome annotation will accelerate. The genome annotation includes the identification of genomic enhancers, likely based on the chromatin features observed at them. The collection of chromatin features usable for enhancer prediction continue to grow, including, for example, the co-regulatory factor cohesin [52, 4] and DNA methylation [53]. The largest collections of chromatin feature data for numerous human cells are produced by the Epigenetic Roadmap data and ENCODE Phase 3 and 4. However, the available data is far from complete. There is a clear need for a large-scale open contest for benchmarking computational and experimental methods for enhancer prediction, also suggested by others [38, 54]. This is still hindered by the lack of sufficient data for different cell types and the lack of a gold-standard set of enhancers and non-enhancers. The set of massive parallel reporter assay (MPRA) validated enhancers could be adopted as the training examples [55, 56], although the set sizes are still rather restricted in the human cells. Moreover, the MPRA approaches have their limitations: the length of the assayed DNA is small, and chromatin environment of the assayed region is not considered in the plasmid-based assay.

## Conclusion

The supervised machine learning methods for the enhancer prediction task may not have adopted the full potential of the probabilistic approach. Therefore, a PRobabilistic Enhancer PRedictIoN tool (PREPRINT) was developed assuming characteristic patterns of chromatin features at enhancers and other genomic regions. PREPRINT included two separate approaches to statistically model the chromatin feature data, the PREPRINT maximum likelihood (ML) and the PREPRINT Bayesian approach. The performances of PREPRINT and the state-of-the-art methods RFECS and ChromHMM were compared on chromatin feature data from the ENCODE first data production phase Tier 1 cell lines K562 and GM12878.

To control the false positive rate of genome-wide enhancer predictions, the prediction threshold for PREPRINT was calibrated. Although the lengths of the PREPRINT enhancer predictions were less sensitive to changes in the prediction threshold than the lengths of the RFECS predictions, the prediction threshold was demonstrated to influence the final number of predicted enhancers and their validation rates. In general, PREPRINT performed comparably to or outperformed the state-of-the-art methods, especially when requiring the methods to predict a larger set of enhancers, i.e., employing a relaxed threshold. Firstly, the PREPRINT Bayesian trained on the data originating from the cell line K562 achieved the best performance on the small test data originating from the GM12878 cell line. Secondly, the proportions of genome-wide predictions lacking a TRF binding site were lower for PREPRINT when requiring a larger set of prediction. In addition, PREPRINT enhancers contained proportionally more enhancers overlapping a small number of different TRF peaks than the predictions obtained by RFECS. Thirdly, when considering the genome-wide predictions, the PREPRINT trained on the K562 data generalised successfully on the GM12878 data, and the Bayesian approach performed sufficiently. In some comparisons, the Bayesian approach attained similar or even superior performance to RFECS. The enhancers predicted by PREPRINT, RFECS and ChromHMM were demonstrated to be highly congruent. Nevertheless, the unique enhancers predicted only by the PREPRINT Bayesian approach possessed robust validation rates, despite varying the prediction threshold. When comparing the two PREPRINT approaches, the performance of PREPRINT Bayesian often exceeded the performance of the ML approach. This implies that a global genome-wide distribution of the scaling parameter (PREPRINT Bayesian) should be adopted, instead of modelling locally the fit of an individual sample, for example, to the characteristic enhancer chromatin feature pattern (PREPRINT ML).

Accurate annotation of the regulatory regions across the human genome is a prerequisite for the interpretation of the findings of regulatory genomics studies, the GWAS studies as well as the clinical studies. In this work, a machine learning tool was developed to predict genomic regulatory enhancers in the human genome based on the chromatin feature data quantified by next-generation sequencing technology. In the future, the next-generation sequencing methodologies will likely transition to standard clinical tests, and in clinical diagnostics, the analysis will be broadened from genotyping the protein-coding genes towards profiling the non-coding DNA.

There are many general problems related to the computational genome-wide enhancer prediction, and future studies are necessary. There is a need to generate a golden standard set of enhancers to develop and benchmark the computational methods. In addition, there are likely enhancer subsets possessing different features and feature patterns. Therefore, clustering as a preprocessing step or semi-supervised approaches should be adopted. The resolution of the data and the prediction precision and accuracy should be increased, for example, to pinpoint the individual enhancers.

## Methods

### PREPRINT Outline

This paper presents a new method called PRobabilistic Enhancer Prediction Tool (PREPRINT). PREPRINT consists of several data processing and analysis steps, which are sketched in Figure 7 as five modules (black rounded boxes). The modules are the data preprocessing (Figure 7**a** and **b**), statistical modelling (Figure 7**c**), computing probabilistic scores to build the final training data matrix (Figure 7**d** and **e**), using the data matrix to train a Support Vector Machine (SVM) classifier with a Gaussian kernel (Figure 7**f**), and finally computing the probabilistic scores for the genome-wide data and employing PREPRINT to obtain genome-wide enhancer predictions (Figure 7**g** and **h**). The five modules are shortly described in the Supplementary Methods section, Additional File 1. A more detailed description of the modules and the steps inside each module are provided in the following sections.

**Figure 7:**
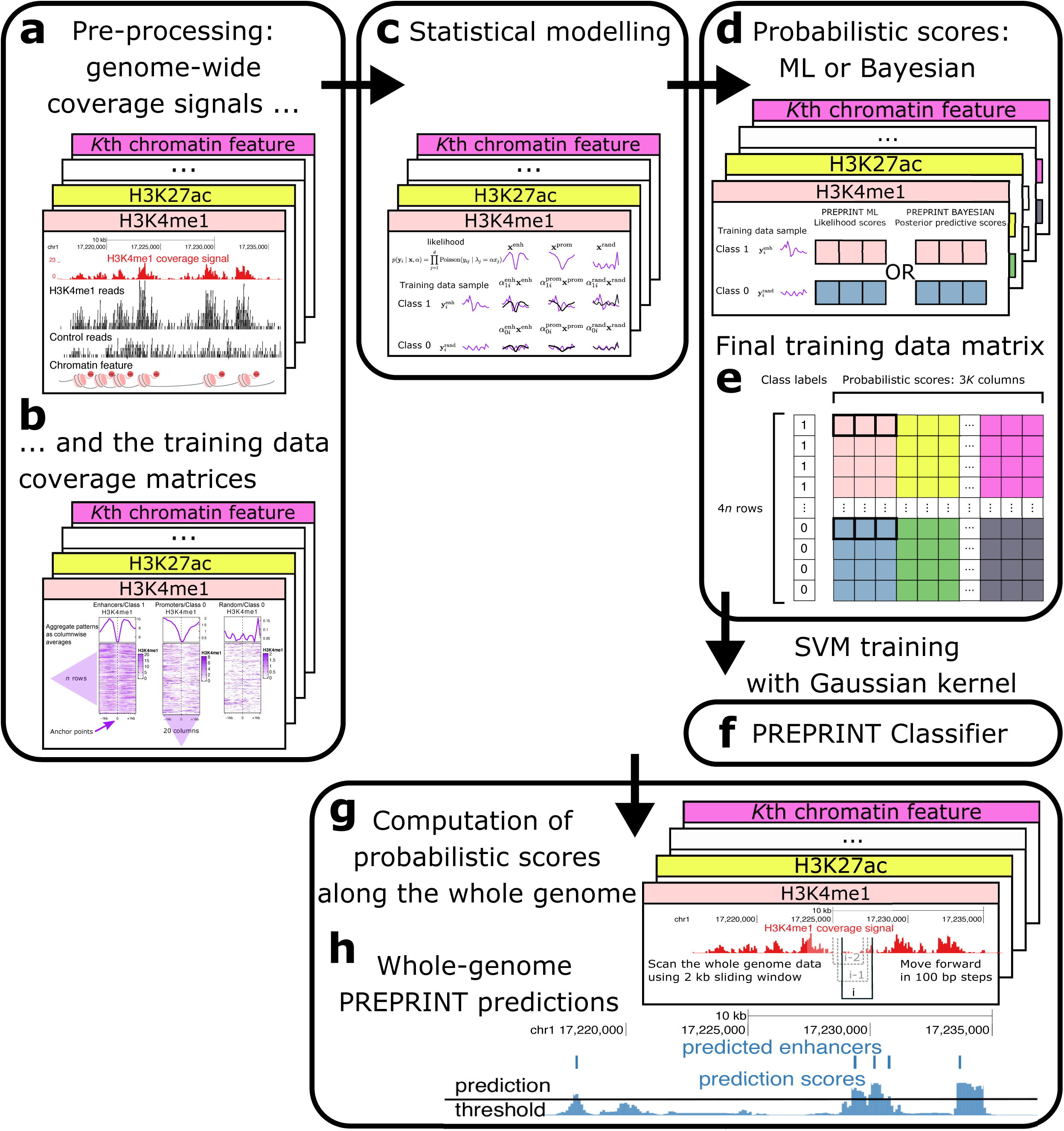
Diagram of the PREPRINT steps. The PREPRINT procedure consists of five modules. In the preprocessing module, the genome-wide coverage signals for *K* chromatin features were first extracted from the short sequencing reads (Step **a**). In the second Step **b**, the training data coverage matrices and the aggregate patterns were obtained. In the statistical modelling module (Step **c**), the individual samples were assumed to follow a Poisson distribution given the scaled aggregate patterns as parameters. In the third module, to quantify the fit of the samples to the aggregate patterns, maximum likelihood (ML) or Bayesian probabilistic scores were computed (Step **d**). The probabilistic scores were collected into the final training data matrix (Step **e**). In the fourth module, an SVM classifier was trained with a Gaussian Kernel (Step **f**). Finally, in the fifth module, the probabilistic scores were computed along the whole genome (Step **g**) and the probabilistic scores were classified by PREPRINT to obtain the enhancer predictions (Step **h**).

### Data

This work adapted the publicly available data from ENCODE Consortium [43]. The data contained the ChIP-seq raw reads from ENCODE/Broad Institute data set for 10 histone modifications, a histone variant H2AZ, and a protein CTCF. The data were downloaded for the myelogenous leukaemia cell line K562 and the lymphoblastoid cell line GM12878. In addition, the RNA polymerase II ChIP-seq data were downloaded from Transcription Factor Binding Sites by ChIP-seq from ENCODE/Stanford/Yale/USC/Harvard data set. Due to the binding of TRFs, enhancers exhibit chromatin openness quantified by DNase I hypersensitivity (DNase I HS) sequencing technique (DNase-seq) [57]. As the DNase-seq data this study utilized the Open Chromatin by DNase I HS data set produced by ENCODE/OpenChrom (Duke University). In addition, the location of nucleosomes at enhancers can be quantified applying micrococcal nuclease digestion followed by sequencing (MNase-seq) [58, 59]. The Nucleosome Position by MNase-seq data set from ENCODE/Stanford/BYU were adopted in this study. The DNase-seq and MNase-seq data were downloaded in the aligned format (bam-format). Data for chromosomes chrY and chrM were excluded from all data. The paths to the downloaded files, their ENCODE Data Coordination Center (DCC) Accession names, Data Coordination Centers (ENCODE DCC), and Gene Expression Omnibus (GEO) sample accession numbers are provided in Additional file 2.

In the preprocessing module of PREPRINT (Figure 7**a**), the ChIP-seq data were processed with the following steps:

1. Raw reads were aligned to the human genome version hg19 with Bowtie 2 [60] (bowtie2-2.3.3.1) adopting the default options.
2. Reads aligned to the exact same locations were considered as Polymerase Chain Reaction (PCR) duplicates [61], and only one of the duplicate reads was retained for the analysis.
3. Possible isogenic replicates were pooled.
4. The fragment lengths for ChIP-sed reads were estimated from the cross-correlation profiles obtained with phantompeakqualtools (spp version 1.14) [62, 63] and R version R-3.3.1.
5. The ChIP-seq reads were shifted by the half of the estimated fragment length with the combination of bedtools2 and samtools, In addition, the MNase-seq reads were shifted by 149*/*2, the half of the length of DNA wrapping around a nucleosome (*∼*149 bps). In contrast, the DNase-seq and control reads were not shifted.
6. The genome-wide coverage signals were generated. When creating the coverage signal, the control coverage was normalised wrt. the ChIP coverage to equalise the library sizes, and the control coverage was subtracted from the ChIP coverage. Data for GM12878 were normalised wrt. the K562 library size. The normalisation was performed as previously described [64, 62, 65].
7. For PREPRINT, the control signal was not subtracted from the DNase-seq and MNase-seq signal. In contrast, RFECS requires the control signal for all data types, as it is adapted to the histone modification ChIP-seq data. Hence, RFECS normalises and subtracts the control from the DNase-seq and MNase-seq signals.
8. For PREPRINT, the coverage was computed in 100 bp bins, the coverage values were rounded to the nearest integer, and the negative values were converted to zero. RFECS also utilised data in 100 bp bins.

### Definition of the training data

This section describes the extraction of the training data matrices in the preprocessing module (Figure 7**b**). In many earlier studies, the binding sites of co-regulatory proteins and histone acetylases, such as CBP or p300, have been used to identify enhancer locations [12, 11, 14, 13, 66]. Therefore, the training and test data enhancer samples were defined as the summits of the 1000 most significant (based on q-values) histone acetylase P300 binding sites. The summit is the location with the highest read pileup within a ChIP-seq peak, often considered as the precise binding location of the TRF. In addition, each P300 binding site summit was required to overlap a DNase I hypersensitivity (HS) peak. As the binding sites of P300, the Transcription Factor ChIP-seq Uniform Peaks from ENCODE/Analysis were adopted in the training cell line K562 (SydhK562P300Iggrab) and test cell line GM12878 (SydhGm12878P300Iggmus). As the DNase I HS peaks, the Open Chromatin data set from ENCODE/OpenChrom(Duke University) was utilised. The distances between the training or test data enhancers and the protein-coding transcription start sites (TSS) from Gencode v27 [67] were required to equal or exceed 2 kb. The enhancers were centred at the p300 peak summits, i.e., the summits were considered as the enhancer anchor points. See Additional File 2 for more details about the origin of the data.

As examples of non-enhancers, promoters were defined as the 1000 GENCODE v27 TSSs that overlapped a DNAse I HS peak and whose distance to any other TSSs nearby exceeded 2 kb. For both cell lines, the 1000 promoters overlapping the most significant DNase-seq peaks (based on the p-values) were selected. The TSSs were used as the anchor points for the promoters. When defining the training data samples, the enhancers and promoters (defined as the 5 kb regions centred at the anchor points) overlapping the ENCODE blacklist regions were excluded. The ENCODE blacklist regions contained repetitive elements, such as *α*-and *β*-satellite repeats, ribosomal and mitochondrial DNA, and some other regions that are listed in the Mappability or Uniqueness of Reference Genome data set [68]. For more details about the ENCODE blacklist regions, see Additional File 2.

For the training and test data enhancers and promoters, 2 kb genomic windows centred at the anchor points were defined, and the coverage matrices of the chromatin features were extracted at 100-bp resolution. Figure 8 illustrates the coverage matrices and the aggregate profiles of the different chromatin features at the training data enhancers. In contrast, Supplementary Figure S1, Additional File 1, illustrates the heatmaps and aggregate profiles at the training data promoters. The promoters were not oriented according to the transcription direction. In addition to promoters, random genomic regions were included as examples of non-enhancers. Thus, 1000 random genomic locations were sampled for both cell lines requiring that the distance between a sampled single bp genomic coordinate and any cell line-specific P300 ChIP-seq peak exceeded 2.5 kb. Moreover, the sampled coordinate was required to have a distance exceeding 2 kb to the nearest TSS. Further, random genomic locations overlapping the ENCODE blacklist regions were removed. These regions were referred to as the pure random regions. The coverage matrices and the aggregate profiles at the pure random regions in the K562 cell line are demonstrated in Supplementary Figure S2, Additional File 1. At the pure random regions, the feature patterns were close to zero, hence the classifier would apparently learn to distinguish the random regions from the enhancers. Therefore, another set of random locations was defined as follows. Firstly, the sum of genome-wide coverage signals excluding the MNase-seq signal was computed in 100 bp bins. The bins with the coverage sum equal to or exceeding 5 were selected. Again, the p300 binding sites, the TSSs and the ENCODE blacklist regions were removed from the selected regions. The selected regions comprised around 4% of the whole genome. The random regions were sampled within the selected regions requiring that the coverage sum in all 100 bp bins of the 2 kb window centred at the anchor point was equal to or exceeded 5. Thus, these random locations acquired some non-zero signal and are referred to as random regions with a signal. The coverage matrices and the aggregate profiles at the 1000 random regions with a signal in the K562 cell line are visualised in Supplementary Figure S3, Additional File 1. To conclude, the final training data contained *n* enhancers, *n* promoters and 2*n* random locations; here *n* = 1000.

**Figure 8:**
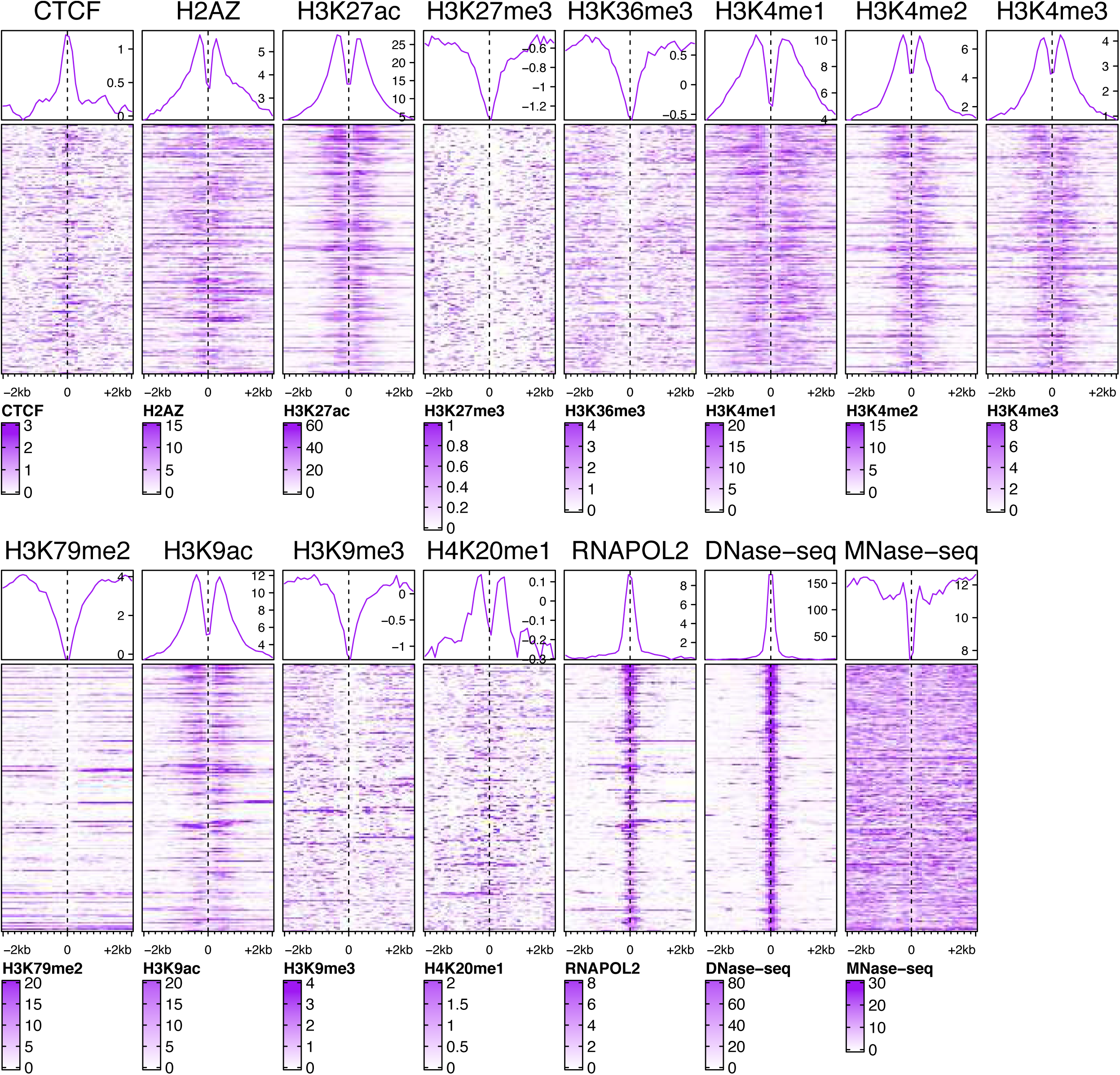
The coverage matrices and the aggregate patterns of 15 different chromatin features at the 1000 enhancer samples. The coverage matrices were visualised as heatmaps together with the aggregate patterns illustrated above the heatmaps. The data originated from cell line K562, and the feature patterns were extracted in a genomic window of length 4 kb centred at the enhancer anchor points indicated by the dashed line and the coordinate 0. The resolution (bin size) of the data was 100 bp.

### Probabilistic modelling

This section describes the second and the third module of the PREPRINT procedure (Figure 7**c, d**, and **e**). The chromatin features were indexed by *k* = 1, …, *K*, where *K* was the total number of chromatin features. The training coverage data for the chromatin feature *k* were represented as a (4*n × d*) matrix **Y**_*k*_ of 4*n* samples and *d* bins, i.e., *d* was the length of the feature patterns. In the training coverage matrix, there were *n* enhancer samples, *n* promoter samples and 2*n* random region samples. The matrix **Y**_*k*_ was further divided into (*n*_enh_ *× d*) training data enhancer matrix 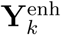, the (*n*_prom_ *× d*) training data promoter matrix 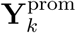, and the (*n*_rand_ *× d*) training data random region matrix 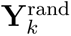. The feature pattern of the chromatin feature *k* for an enhancer sample *i* was denoted as 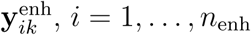. The elements of the feature pattern vector 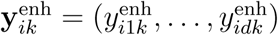 indexed by *j* = 1, …, *d* were assumed to follow a Poisson distribution. Further assuming the conditional independence of the feature pattern elements, the likelihood of 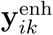 was

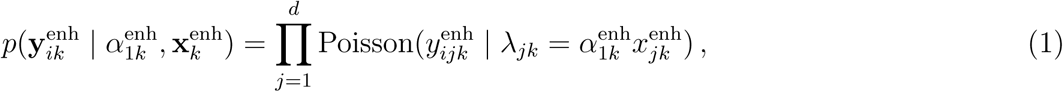

where *λ*_*jk*_ denotes the rate parameter of the *j*th feature pattern element. The parameter *λ*_*jk*_ was an auxiliary variable and a product of the aggregate pattern 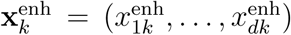 and a scaling parameter 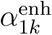. The subindex 1 in 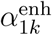 referred to the class of sample 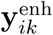, namely 1 (see also Figure 7**c**). In this model, the enhancer aggregate pattern 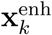 captured the characteristic pattern of the chromatin feature *k* at enhancers, and the scaling parameter 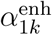 was shared between the elements of the feature pattern vector 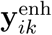. The scaling parameter 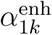 modelled the statistical variation of the feature pattern elements at enhancers resulting from the sources listed in the Background Section. Nevertheless, the variability in the feature pattern elements were assumed to originate from a local source. In other words, the variation was shared by the elements of the feature pattern vector. A natural choice to model the scaling parameter 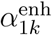 was the Gamma distribution with the hyperparameters *a*_0*k*_ and *b*_0*k*_, the conjugate distribution of the Poisson distribution

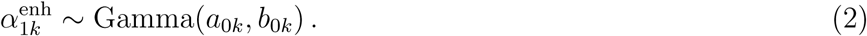

The parameters of the model were estimated from the data as follows: Firstly, the variable 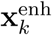 was estimated as the training data enhancer average coverage, i.e., the aggregate pattern. The elements of the aggregate pattern were computed as

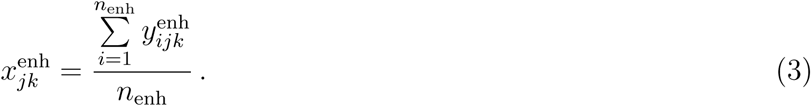

Secondly, the Gamma distribution of the scaling parameter 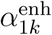 was learned by first obtaining the maximum likelihood (ML) estimates of 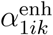 for each training data enhancer sample. The likelihood in Equation 1 was maximized to obtain the ML estimates

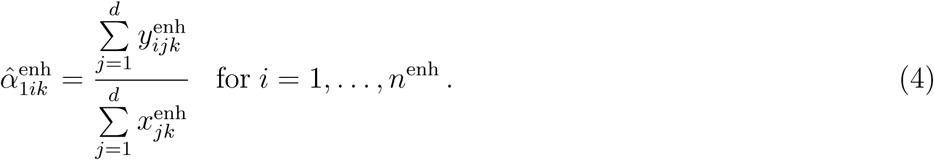

Next, a Gamma distribution was fitted to 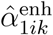 values to obtain the empirical estimates for the hyperparameters *a*_0*k*_ and *b*_0*k*_. Finally, to quantify the fit of any training data sample *i* = 1, …, 4*n* to the enhancer aggregate pattern, a posterior predictive score was computed as

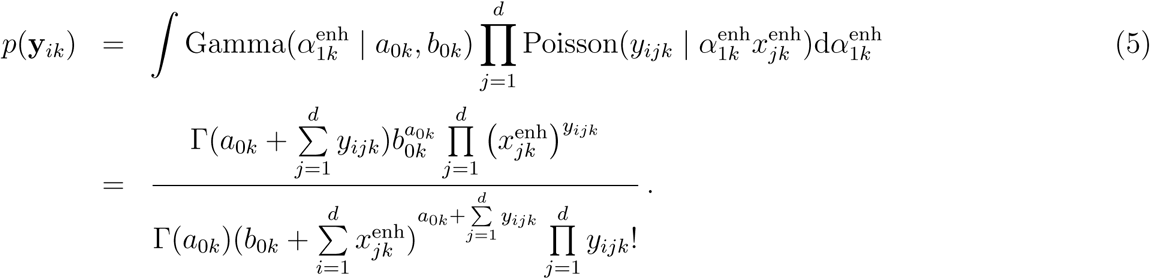

The posterior predictive scores are referred to as probabilistic scores. As the individual training data samples were considered independent, the posterior predictive scores could be also derived for the individual samples as provided in Equation 5; there were no product over all the samples. Equation 5 resembled the Gamma-Poisson compound distribution, equivalent to the negative binomial distribution. However, the Equation 5 could not be simplified into the negative binomial distribution due to the product over *d* adjacent bins and the product *λ*_*jk*_ = *α*_*k*_*x*_*jk*_ (sub- and superindices omitted for clarity); this removed the conjugacy between Gamma and Poisson distributions. The computation of the probabilistic score to quantify the fit between each training data sample and the enhancer aggregate patterns of *K* chromatin features resulted in a matrix of size 4*n × K*.

The genomic regions with patterns of chromatin features resembling the aggregate patterns of enhancer while lacking the characteristic patterns of promoters and random regions should receive a high prediction score. Therefore, in addition to quantifying the fit of the individual sample to the enhancer aggregate pattern, we computed the fit of the sample to the non-enhancer aggregate pattern. The elements of the promoter aggregate pattern were computed as

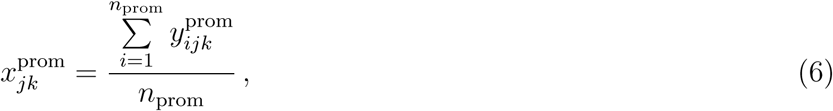

and the ML estimates of 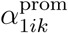 (see Figure 7**c**) were computed as

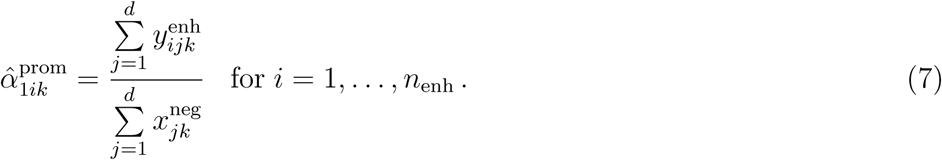

Notably, the parameter 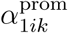 scales the promoter aggregate profile to fit the training data enhancer sample. Again, a Gamma distribution was fitted to the 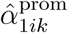 values to obtain the (empirical) estimates for the hyper-parameters 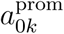 and 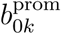. Then the probabilistic scores of the samples were computed again using Equation 5. However, instead of integrating over 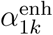, the integral was over 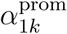. Furthermore, the scaling parameters 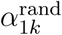 (see Figure 7**c**) corresponding to the random aggregate patterns were estimated similarly as in Equation 7. The quantification of the fit of any training data sample to the random aggregate pattern were computed again using the Equation 5, this time the integral was over 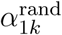. Finally, the fit of any training data sample to the aggregate pattern of enhancer, promoter and random regions resulted in three posterior predictive scores (Figure 7**d**), and for all *K* chromatin features, the final training data vector consisted of 3*K* elements (see Figure 7**e**). This approach is referred to as PREPRINT Bayesian, due to the computation of the posterior predictive scores.

For comparison to the PREPRINT Bayesian approach, a simplified model was considered where the scaling parameters 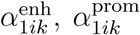, and 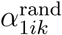 were estimated individually for each training and test data enhancer using Equations 4 and 7. In addition, for a non-enhancer (class 0) sample **y**_*ik*_, the scaling parameters were denoted as 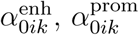, and 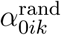 (see Figure 7**c**). The ML estimates for these were obtained, for example, as

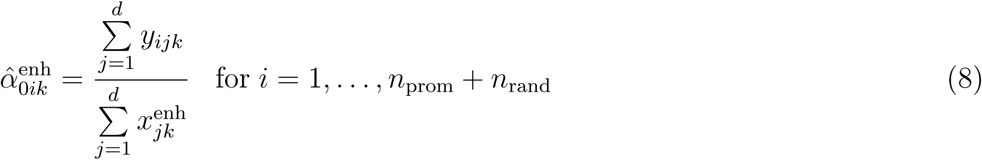

and

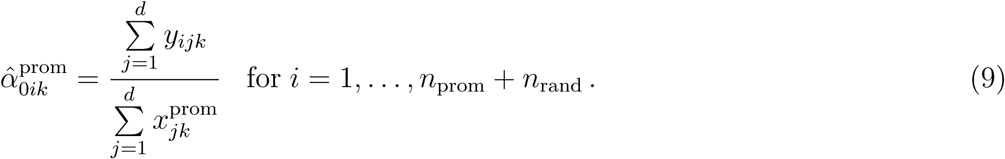

To quantify the fit of any sample pattern to the enhancer, promoter and random aggregate pattern, the likelihood values were computed as the probabilistic scores. For the training and test data enhancers (class 1) the likelihood was computed, for example, as

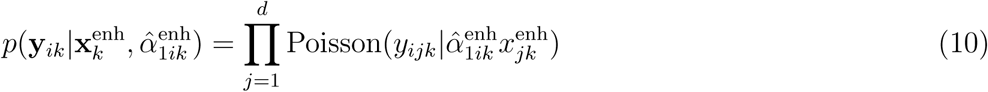

and

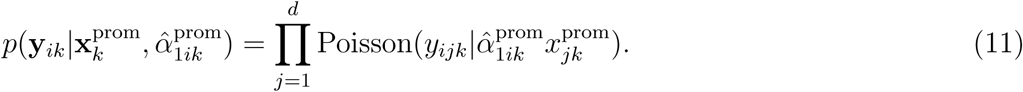

The likelihood values for the training and test data non-enhancers (class 0) samples were computed, for example, as

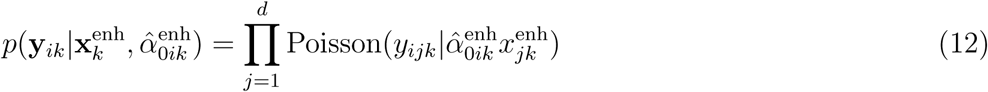

and

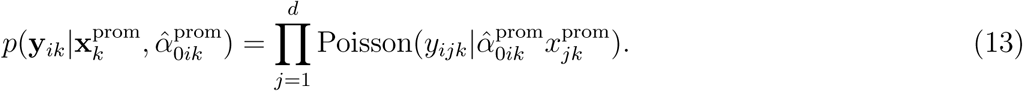

For each sample, three separate likelihood scores were computed corresponding to the fit between the sample and the enhancer, promoter and random region aggregate patterns (see Figure 7**d**). This simpler approach is referred to as PREPRINT maximum likelihood (ML). In the PREPRINT ML, a sample specific fixed scaling parameter *α*_*ik*_ was estimated for each sample pattern individually. The ML estimate for *α*_*ik*_ scaled the aggregate pattern to fit the individual sample pattern, and the fit was quantified by the likelihood score. In contrast, the PREPRINT Bayesian approach involved global Gamma distributions for the scaling parameters associated to the enhancer, promoter and random region aggregate patterns. In other words, in the PREPRINT Bayesian approach, the probabilistic score in Equation 5 was computed as the likelihood integrated over the distribution of the scaling parameter *α*_1*k*_.

### Classifier training and cross-validation, performance on the test set

This section describes the fourth module in the PREPRINT procedure (Figure 7**f**). A support vector machine (SVM) classifier implemented in libSVM version 3.22 [69] was trained on the final training data matrix presented in Figure 7**e**. The training data originated from cell line K562, and the SVM utilised a Gaussian kernel. The hyperparameters of the SVM classifier were estimated and the classifier performance was evaluated by a nested cross-validation (CV). In the nested CV, the outer cross-validation assessed the performance of the model and the inner optimised the hyperparameters, namely the SVM misclassification penalty *C* and the Gaussian kernel width *γ*. The hyperparameter optimisation was performed by a grid-search with values *C* = 2^*-*5^, 2^*-*4.5^, …, 2^25^ and *γ* = 2^*-*25^, 2^*-*24.5^, …, 2^10^. For both the outer and inner cross-validation, a 5-fold CV was adopted. The method performance for cell line K562 was evaluated by concatenating the predictions obtained from separate cross-validation rounds and computing the area under the receiver operating characteristics curve (AUC). The final PREPRINT classifier was trained on the whole training data matrix from the K562 data, and again a 5-fold CV was performed to optimise the SVM hyperparameters. The final classifier predicted enhancers on the GM12878 test data.

The classification performance of PREPRINT was compared to RFECS, a state-of-the-art supervised enhancer prediction method. For RFECS, the CV within K562 training data was not performed; the AUC values would have likely achieved values close to 1. The performance RFECS trained on K562 data was evaluated on the GM12878 test data as follows. Firstly, the RFECS predicted enhancers genome-wide in cell line GM12878. The genome-wide RFECS predictions closest to the test regions were selected. The performance on the test set was again evaluated by the AUC values.

### Genome-wide enhancer predictions and their validation

The PREPRINT and RFECS classifiers trained on the K562 training data predicted enhancers genome-wide in both cell lines, K562 and GM12878. Before studying the properties of the predicted enhancers and validating the results, the training data enhancers and promoters were removed from the final genome-wide predictions. In addition, any obscure genomic regions were removed from the genome-wide predictions. The removed regions included predictions (defined as 2 kb genomic windows) overlapping at least 1 bp with the ENCODE blacklist regions as well as the K562 cell line predictions with a distance equal or smaller than 1 kb to any training data enhancer. To remove promoters from the set of enhancer predictions, we excluded the predictions whose middle base had a distance equal or smaller than 2 kb to any GENCODE v27 transcription start site (TSS).

The genome-wide enhancer predictions were validated by inspecting the overlap between the predicted enhancers and the histone acetyltransferase (p300) binding sites (SydhK562P300Iggrab and SydhGm12878P300Iggmus ChIP-seq peak sets). In addition, a large set of TRF binding sites from the Transcription factor ChIP-seq Uniform peaks produced by ENCODE were utilised for validation. The peaks for RNA Pol II, CTCF, CREB-binding protein (CBP) and p300 were excluded from the Uniform peak set resulting in peaks for 111 and 76 individual TRFs for cell lines K562 and GM12878, respectively. For more details about the validation data, see Additional File 2. An enhancer was validated if the 2 kb prediction window overlapped with at least 1 bp of at least 1 TRF ChIP-seq peak. The ChromHMM enhancer clusters contained regions with varying sizes; these were validated similarly by requiring at least 1 bp overlap between the variable size ChromHMM enhancer region and at least 1 TRF ChIP-seq peak. When considering the number of overlapping TRF ChIP-seq peaks at a given enhancer prediction, the peaks for the individual TRFs were not required to overlap, an overlap was required only between the 2 kb prediction window and a single TRF ChIP-seq peak.

## List of abbreviations

AUC: Area Under the receiver operating characteristics Curve
bp: Base pair
CBP: CREB-binding-protein
ChIP-seq: Chromatin Immunoprecipitation coupled with Sequencing
CNV: Copy number variation CRE-seq Cis-regulatory element analysis by sequencing reporter assay
CTCF: CCCTC-binding factor
CV: Cross-validation
DNase I HS: DNase I hypersensitivity
DNase-seq: DNase I hypersensitive sites sequencing
EM: Expectation-Maximisation
ENCODE: Encyclopedia of DNA Elements
FPR: False positive rate
GENCODE: The reference genome annotation for human and other species
GM12878: Lymphoblastoid cell line
GWAS: Genome-wide association study
H2AZ: Histone variant
H3K27ac: Histone 3 lysine 27 acetylation
H3K27me3: Histone 3 lysine 27 trimethylation
H3K36me3: Histone 3 lysine 36 trimethylation
H3K4me3: Histone 3 lysine 4 trimethylation
H3K79me2: Histone 3 lysine 79 dimethylation
H3K9ac: Histone 3 lysine 9 acetylation
H4K20me1: Histone 4 lysine 20 monomethylation
K562: Myelogenous leukaemia cell line
kb: Kilobase
ML: Maximum likelihood
MNase-seq: Micrococcal nuclease digestion followed by sequencing
MPRA: Massive parallel reporter assay
NGS: Next-generation sequencing p300: Histone acetyltransferase
PCR: Polymerase Chain Reaction
PREPRINT: Probabilistic enhancer prediction tool
RFECS: Random Forest-based Enhancer Identi-fication from Chromatin States
RNA Pol II: RNA polymerase II
ROC: Receiver operating characteristics
SNP: Single nucleotide polymorphism
SVM: Support vector machine
TF: Transcription factor
TRF: Transcription related factor
TSS: Transcription start site

## Declarations

### Ethics approval and consent to participate

Not applicable.

### Consent for publication

Not applicable.

### Availability of data and material

The download links to the original data analysed in this study are included in the supplementary information files of this published article. The data used in this work and the links to download the data are listed in the Additional File 2. The PREPRINT package and the codes for the data preprocessing steps are available in GitHub https://github.com/MariaOsmala/preprint. The data and enhancer predictions are stored as a UCSC Genome Browser track hubs, links to the track hubs are provided in GitHub.

## Ethics approval and consent to participate

Not applicable.

## Author’s contributions

MO and HL developed the machine learning methods. MO compiled the data sets, implemented the method, performed the computational analysis and wrote the manuscript. HL participated in the evaluations of the findings and revision of the manuscript. Both authors read and approved the final manuscript.

## Acknowledgements

We thank our colleagues in the Lähdesmäki laboratory for discussions and feedback regarding this manuscript. We thank Dr. Gökçen Eraslan and Dr. Nisha Rajagopal providing support to execute the RFECS experiments. The calculations presented above were performed using computer resources within the Aalto University School of Science “Science-IT” project.

## Additional Files

### Additional file 1 — Supplementary Methods, Figures and Tables

This file contains the Supplementary Methods section, the Supplementary **Figures S1–S8**, and the Supplementary **Tables S1–S2**. PDF of size 94000 kB.

### Additional file 2 — The origin of data

The links to the downloaded files, their ENCODE Data Coordination Center (DCC) Accession names, Data Coordination Centers (ENCODE DCC), and Gene Expression Omnibus (GEO) sample accession numbers. Excel (.xlsx) file of size 34.4 kB.

### Additional file 3

The full genome browser example figure of the K562 cell line data. PDF of size 199 kB.

### Additional file 4

The full genome browser example figure of the GM12878 cell line GM12878. PDF of size 217 kB.

## References

[1] Karnuta JM, Scacheri PC. Enhancers: bridging the gap between gene control and human disease. Human molecular genetics. 2018;27(R2):R219–R227.

[2] Corradin O, Scacheri PC. Enhancer variants: Evaluating functions in common disease. Genome Medicine. 2014;6(10):85.

[3] Smith E, Shilatifard A. Enhancer biology and enhanceropathies. Nature Structural and Molecular Biology. 2014;21(3):210–219.

[4] Shlyueva D, Stampfel G, Stark A. Transcriptional enhancers: From properties to genome-wide predictions. Nature Reviews Genetics. 2014;15(4):272–286.

[5] Long HK, Prescott SL, Wysocka J. Ever-Changing Landscapes: Transcriptional Enhancers in Development and Evolution. Cell. 2016;167(5):1170–1187.

[6] Rickels R, Shilatifard A. Enhancer Logic and Mechanics in Development and Disease. Trends in Cell Biology. 2018;28(8):608–630.

[7] Banerji J, Rusconi S, Schaffner W. Expression of a *β*-globin gene is enhanced by remote SV40 DNA sequences. Cell. 1981;27(2):299–308.

[8] Moreau P, Hen R, Wasylyk B, Everett R, Gaub MP, Chambon P. The SV40 72 base repair repeat has a striking effect on gene expression both in SV40 and other chimeric recombinants. Nucleic Acids Research. 1981;9(22):6047–6068.

[9] Johnson DS, Mortazavi A, Myers RM, Wold B. Genome-wide mapping of in vivo protein-DNA interactions. Science. 2007;316(5830):1497–1502.

[10] Robertson G, Hirst M, Bainbridge M, Bilenky M, Zhao Y, Zeng T, et al. Genome-wide profiles of STAT1 DNA association using chromatin immunoprecipitation and massively parallel sequencing. Nature Methods. 2007;4(8):651–657.

[11] Gilbert J, Drenkow J, Bell I, Zhao X, Srinivasan KG, Sung WK, et al. Identification and analysis of functional elements in 1% of the human genome by the ENCODE pilot project. Nature. 2007;447(7146):799–816.

[12] Heintzman ND, Stuart RK, Hon G, Fu Y, Ching CW, Hawkins RD, et al. Distinct and predictive chromatin signatures of transcriptional promoters and enhancers in the human genome. Nature genetics. 2007;39(3):311.

[13] Heintzman ND, Hon GC, Hawkins RD, Kheradpour P, Stark A, Harp LF, et al. Histone modifications at human enhancers reflect global cell-type-specific gene expression. Nature. 2009;459(7243):108–112.

[14] Visel A, Blow MJ, Li Z, Zhang T, Akiyama JA, Holt A, et al. ChIP-seq accurately predicts tissue-specific activity of enhancers. Nature. 2009;457(7231):854–858.

[15] Rada-Iglesias A, Bajpai R, Swigut T, Brugmann SA, Flynn RA, Wysocka J. A unique chromatin signature uncovers early developmental enhancers in humans. Nature. 2011;470(7333):279–285.

[16] Creyghton MP, Cheng AW, Welstead GG, Kooistra T, Carey BW, Steine EJ, et al. Histone H3K27ac separates active from poised enhancers and predicts developmental state. Proceedings of the National Academy of Sciences of the United States of America. 2010;107(50):21931–21936.

[17] Spitz F, Furlong EEM. Transcription factors: From enhancer binding to developmental control. Nature Reviews Genetics. 2012;13(9):613–626.

[18] Zabidi MA, Stark A. Regulatory Enhancer–Core-Promoter Communication via Transcription Factors and Cofactors. Trends in Genetics. 2016;32(12):801–814.

[19] Elnitski L, Jin VX, Farnham PJ, Jones SJM. Locating mammalian transcription factor binding sites: A survey of computational and experimental techniques. Genome Research. 2006;16(12):1455–1464.

[20] Su J, Teichmann SA, Down TA. Assessing computational methods of cis-regulatory module prediction. PLoS Computational Biology. 2010;6(12):e1001020.

[21] Hardison RC, Taylor J. Genomic approaches towards finding cis-regulatory modules in animals. Nature Reviews Genetics. 2012;13(7):469–483.

[22] Sheffield NC, Furey TS. Identifying and characterizing regulatory sequences in the human genome with chromatin accessibility assays. Genes. 2012;3(4):651–670.

[23] Kleftogiannis D, Kalnis P, Bajic VB. Progress and challenges in bioinformatics approaches for enhancer identification. Briefings in Bioinformatics. 2016;17(6):967–979.

[24] Lim LWK, Chung HH, Chong YL, Lee NK. A survey of recently emerged genome-wide computational enhancer predictor tools. Computational Biology and Chemistry. 2018;74:132–141.

[25] Ernst J, Kellis M. Discovery and characterization of chromatin states for systematic annotation of the human genome. Nature Biotech. 2010;28:817–825.

[26] Ernst J, Kheradpour P, Mikkelsen TS, Shoresh N, Ward LD, Epstein CB, et al. Mapping and analysis of chromatin state dynamics in nine human cell types. Nature. 2011;473:43–49.

[27] Rajagopal N, Xie W, Li Y, Wagner U, Wang W, Stamatoyannopoulos J, et al. RFECS: A Random-Forest Based Algorithm for Enhancer Identification from Chromatin State. PLoS Computational Biology. 2013;9(3):e1002968.

[28] Won KJ, Chepelev I, Ren B, Wang W. Prediction of regulatory elements in mammalian genomes using chromatin signatures. BMC Bioinformatics. 2008;9:547.

[29] Firpi HA, Ucar D, Tan K. Discover regulatory DNA elements using chromatin signatures and artificial neural network. Bioinformatics. 2010;26(13):1579–1586.

[30] Fernández M, Miranda-Saavedra D. Genome-wide enhancer prediction from epigenetic signatures using genetic algorithm-optimized support vector machines. Nucleic Acids Research. 2012;40(10):e77.

[31] Marioni JC, Mason CE, Mane SM, Stephens M, Gilad Y. RNA-seq: An assessment of technical reproducibility and comparison with gene expression arrays. Genome Research. 2008;18(9):1509–1517.

[32] Zhang Y, Liu T, Meyer CA, Eeckhoute J, Johnson DS, Bernstein BE, et al. Model-based analysis of ChIP-Seq (MACS). Genome biology. 2008;9(9):R137.

[33] Anders S, Huber W. Differential expression analysis for sequence count data. Genome Biology. 2010;11(10):R106–R106.

[34] Robinson MD, Oshlack A. A scaling normalization method for differential expression analysis of RNA-seq data. Genome Biology. 2010;11(3):R25.

[35] Spyrou C, Stark R, Lynch A, Tavare S. BayesPeak: Bayesian analysis of ChIP-seq data. BMC Bioinformatics. 2009;10(1):299.

[36] Hashimoto TB, Edwards MD, Gifford DK. Universal Count Correction for High-Throughput Sequencing. PLoS Computational Biology. 2014;10(3):e1003494.

[37] Fishilevich S, Nudel R, Rappaport N, Hadar R, Plaschkes I, Iny Stein T, et al. GeneHancer: genome-wide integration of enhancers and target genes in GeneCards. Database. 2017 Jan 1;2017.

[38] Ho EYK, Cao Q, Gu M, Chan RWL, Wu Q, Gerstein M, et al. Shaping the nebulous enhancer in the era of high-throughput assays and genome editing. Briefings in Bioinformatics. 2019 Mar 20;2019, bbz030.

[39] Buecker C, Wysocka J. Enhancers as information integration hubs in development: lessons from genomics. Trends in Genetics. 2012;28(6):276–284.

[40] Xie D, Boyle AP, Wu L, Zhai J, Kawli T, Snyder M. Dynamic trans-acting factor colocalization in human cells. Cell. 2013;155(3):713–724.

[41] Dogan N, Wu W, Morrissey CS, Chen KB, Stonestrom A, Long M, et al. Occupancy by key transcription factors is a more accurate predictor of enhancer activity than histone modifications or chromatin accessibility. Epigenetics and Chromatin. 2015;8(1):16.

[42] Zacher B, Michel M, Schwalb B, Cramer P, Tresch A, Gagneur J. Accurate promoter and enhancer identification in 127 ENCODE and roadmap epigenomics cell types and tissues by GenoSTAN. PLoS ONE. 2017;12(1):e0169249.

[43] Dunham I, Kundaje A, Aldred SF, Collins PJ, Davis CA, Doyle F, et al. An integrated encyclopedia of DNA elements in the human genome. Nature. 2012;489(7414):57–74.

[44] Lambert SA, Jolma A, Campitelli LF, Das PK, Yin Y, Albu M, et al. The Human Transcription Factors. Cell. 2018;172(4):650–665.

[45] Teytelman L, Thurtle DM, Rine J, van Oudenaarden A. Highly expressed loci are vulnerable to misleading ChIP localization of multiple unrelated proteins. Proceedings of the National Academy of Sciences. 2013;110(46):18602–18607.

[46] Wreczycka K, Franke V, Uyar B, Wurmus R, Bulut S, Tursun B, et al. HOT or not: examining the basis of high-occupancy target regions. Nucleic acids research. 2019;47(11):5735–5745.

[47] Strackee J, van der Gon JJD. The frequency distribution of the difference between two Poisson variates. Statistica Neerlandica. 1962;16(1):17–23.

[48] Song Q, Smith AD. Identifying dispersed epigenomic domains from ChIP-Seq data. Bioinformatics. 2011;27(6):870–871.

[49] Kundaje A, Kyriazopoulou-Panagiotopoulou S, Libbrecht M, Smith CL, Raha D, Winters EE, et al. Ubiquitous heterogeneity and asymmetry of the chromatin environment at regulatory elements. Genome Research. 2012;22(9):1735–1747.

[50] Nielsen FGG, Markus KG, Friborg RM, Favrholdt LM, Stunnenberg HG, Huynen M. CATCHprofiles: Clustering and alignment tool for chip profiles. PLoS ONE. 2012;7(1):e28272.

[51] Nair NU, Kumar S, Moret BME, Bucher P. Probabilistic partitioning methods to find significant patterns in ChIP-Seq data. Bioinformatics. 2014;30(17):2406–2413.

[52] Calo E, Wysocka J. Modification of Enhancer Chromatin: What, How, and Why? Molecular Cell. 2013;49(5):825–837.

[53] Fleischer T, Tekpli X, Mathelier A, Wang S, Nebdal D, Dhakal HP, et al. DNA methylation at enhancers identifies distinct breast cancer lineages. Nature Communications. 2017;8(1):1379.

[54] Li Y, Shi W, Wasserman WW. Genome-wide prediction of cis-regulatory regions using supervised deep learning methods. BMC Bioinformatics. 2018;19(1).

[55] Kwasnieski JC, Fiore C, Chaudhari HG, Cohen BA. High-throughput functional testing of ENCODE segmentation predictions. Genome Research. 2014;24(10):1595–1602.

[56] Kheradpour P, Ernst J, Melnikov A, Rogov P, Wang L, Zhang X, et al. Systematic dissection of regulatory motifs in 2000 predicted human enhancers using a massively parallel reporter assay. Genome Research. 2013;23(5):800–811.

[57] Thurman RE, Rynes E. The accessible chromatin landscape of the human genome. Nature. 2012;489(7414):75.

[58] Cui K, Zhao K. Genome-wide approaches to determining nucleosome occupancy in metazoans using MNase-Seq. In: Morse RH, editor. Methods in Molecular Biology. vol. 833. Humana Press; 2012. p. 413–419.

[59] Schones DE, Cui K, Cuddapah S, Roh TY, Barski A, Wang Z, et al. Dynamic Regulation of Nucleosome Positioning in the Human Genome. Cell. 2008;132(5):887–898.

[60] Langmead B, Salzberg SL. Fast gapped-read alignment with Bowtie 2. Nature Methods. 2012;9(4):357–359.

[61] Marx V. How to deduplicate PCR. Nature Methods. 2017;14(5):473–476.

[62] Landt SG, Marinov GK, Kundaje A, Kheradpour P, Pauli F, Batzoglou S, et al. ChIP-seq guidelines and practices of the ENCODE and modENCODE consortia. Genome Research. 2012;22(9):1813–1831.

[63] Kharchenko PV, Tolstorukov MY, Park PJ. Design and analysis of ChIP-seq experiments for DNA-binding proteins. Nature Biotechnology. 2008;26(12):1351–1359.

[64] Li Q, Brown JB, Huang H, Bickel PJ. Measuring reproducibility of high-throughput experiments. Annals of Applied Statistics. 2011;5(3):1752–1779.

[65] Le Martelot G, Canella D, Symul L, Migliavacca E, Gilardi F, Liechti R, et al. Genome-Wide RNA Polymerase II Profiles and RNA Accumulation Reveal Kinetics of Transcription and Associated Epigenetic Changes During Diurnal Cycles. PLoS Biology. 2012;10(11):e1001442.

[66] Kim TK, Hemberg M, Gray JM, Costa AM, Bear DM, Wu J, et al. Widespread transcription at neuronal activity-regulated enhancers. Nature. 2010;465(7295):182–187.

[67] Frankish A, Diekhans M, Ferreira AM, Johnson R, Jungreis I, Loveland J, et al. GENCODE reference annotation for the human and mouse genomes. Nucleic Acids Research. 2019;47(D1):D766–D773.

[68] Carroll TS, Liang Z, Salama R, Stark R, de Santiago I. Impact of artifact removal on ChIP quality metrics in ChIP-seq and ChIP-exo data. Frontiers in Genetics. 2014;5:75.

[69] Chang CC, Lin CJ. LIBSVM: A Library for support vector machines. ACM Transactions on Intelligent Systems and Technology. 2011;2(3):1–27.

