## Supplementary figures and images for "Enhancer prediction in the human genome by probabilistic modelling of the chromatin feature patterns"

### Additional File 3

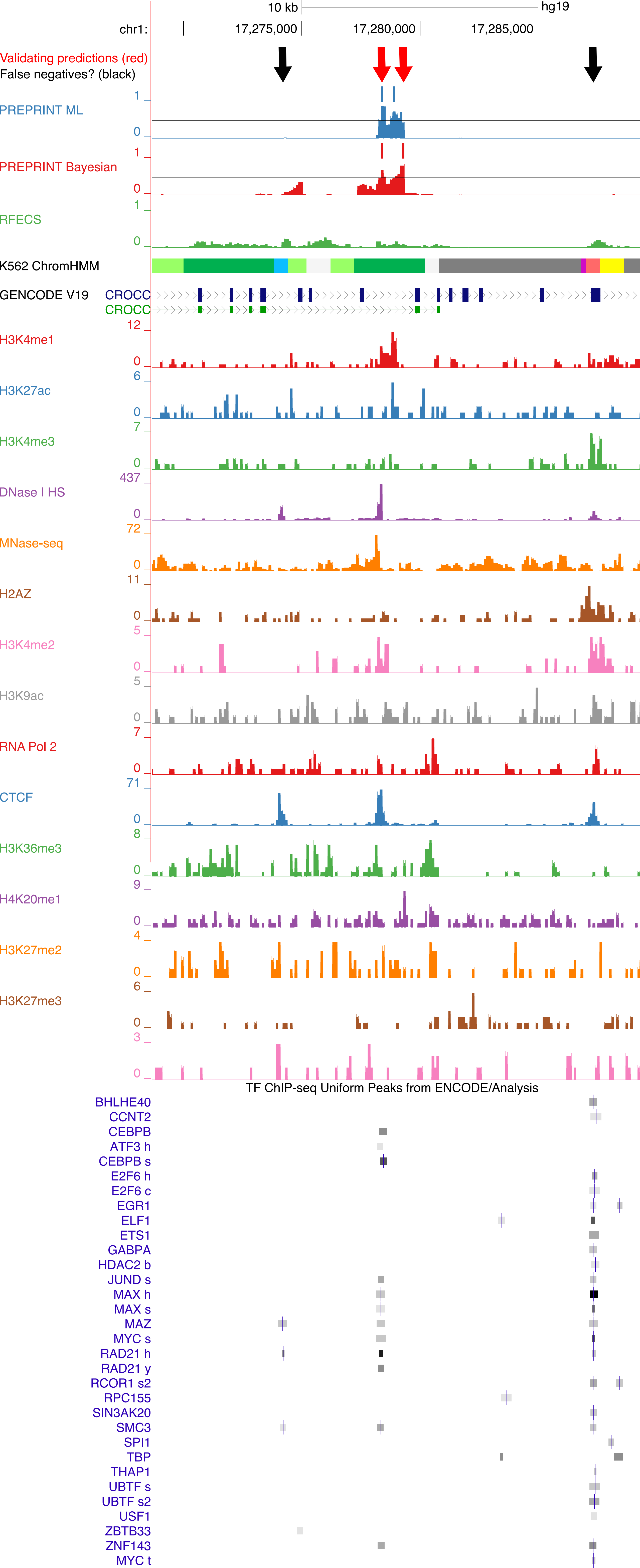

### Additional File 4

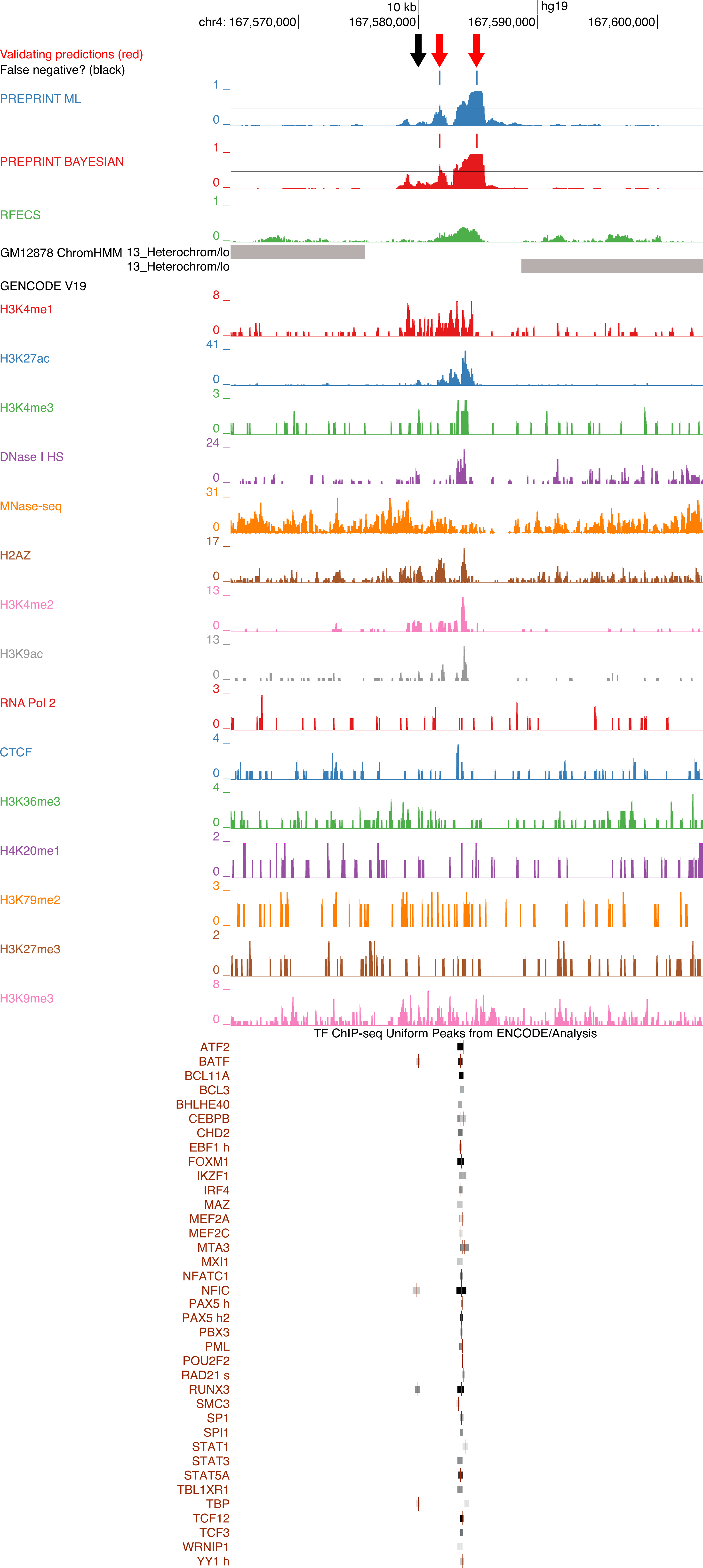
