## Additional File 1 for "Enhancer prediction in the human genome by probabilistic modelling of the chromatin feature patterns"

### List of Figures

S1 The coverage matrices and the aggregate patterns of the 15 chromatin features at the 1000 promoter samples. The coverage matrices were visualised as heatmaps together with the aggregate patterns illustrated above the heatmaps. The data originated from the K562 cell line, and the feature patterns were extracted in a genomic window of length 4 kb centred at the promoter anchor points (TSS) indicated by the dashed line and the coordinate 0. The resolution (bin size) of the data was 100 bp. The promoters are unoriented, i.e., the direction of transcription from TSS is not utilised to direct the coverage patterns. . . . . . . . . . . . . . . . . . . . . . . . . . . . . . . . . . . . 4

S2 The coverage matrices and the aggregate patterns of the 15 chromatin features at the 1000 pure random samples. The coverage matrices were visualised as heatmaps together with the aggregate patterns illustrated above the heatmaps. The data originated from the K562 cell line, and the feature patterns were extracted in a genomic window of length 4 kb centred at the anchor points of the random regions, indicated by the dashed line and the coordinate 0. The resolution (bin size) of the data was 100 bp. . . . . . . . . . . . . . 5

S3 The coverage matrices and the aggregate patterns of the 15 chromatin features at the 1000 random regions with a signal. The coverage matrices were visualised as heatmaps together with the aggregate patterns illustrated above the heatmaps. The data originated from the K562 cell line, and the feature patterns were extracted in a genomic window of length 4 kb centred at the anchor points of the random regions, indicated by the dashed line and the coordinate 0. The resolution (bin size) of the data was 100 bp. . . . . . . 6

S4 The normalised frequencies of the enhancer lengths. The enhancers were predicted in the GM12878 cell line by PREPRINT and RFECS with the thresholds of 0.5 and 0.75. For each method and threshold, the frequencies were divided by the total number of regions predicted as enhancers. The regions were formed by combining the subsequent enhancer predictions into a single region. . . . . . . . . . . . . . . . . . . . . . . . . . . . . . . . . . . 8

S5 The proportions of the genome-wide enhancer predictions overlapping the varying number of TRF ChIP-seq peaks in the GM12878 cell line. The number of enhancers in each comparison are shown above the figure. In comparison **a**, the number of enhancers was the number of enhancers predicted by RFECS with the threshold of 0.25, and in comparison **b**, the number of enhancers was the minimum number of enhancers predicted by PREPRINT methods with their 1% FPR thresholds estimated on the K562 CV data. . . . . . . . . . . 9

S6 The validation rate of the genome-wide enhancer predictions obtained by the different methods and thresholds in the GM12878 cell line. An enhancer prediction was validated if the 2 kb prediction window overlapped with at least 1 bp of at least one TRF ChIP-seq peak. . . . . . . . . . . . . . . . . 10

S7 The unique and overlapping genome-wide enhancer predictions obtained by different methods in the GM12878 cell line. In comparisons **a** and **c**, the PREPRINT predictions were obtained by the ML approach, and in comparisons **b** and **d**, the predictions were obtained by the Bayesian approach. The overlap between the PREPRINT, RFECS and ChromHMM predictions were quantified as the number of enhancers. In each comparison, the number of enhancers predicted by PREPRINT and RFECS was equal. The numbers were: **a** 17746, **b** 17746, **c** 49699, and **d** 49699. Inside every region or intersection, the number of enhancers in the given set is indicated together with the percentage of validated enhancers in the set. The areas of the intersection sets are not proportional to the number of overlapping regions due to the asymmetry of overlaps. . . . . . . . . . . . . . . . . . . . . . . . . . . . 11
S8 The unique and overlapping genome-wide enhancer predictions obtained by PREPRINT and RFECS in the K562 cell line (**a** and **b**) and in the GM12878 cell line (**c** and **d**). In each comparison, the number of enhancers predicted by PREPRINT and RFECS was equal. The numbers were: **a** the minimum number of enhancers predicted by PREPRINT with the 1 % FPR threshold in the K562 cell line (15531), **b** the number of enhancers predicted by RFECS with the threshold of 0.25 in the K562 cell line (35089), **c** the minimum number of enhancers predicted by PREPRINT with the 1 % FPR threshold in the GM12878 cell line (22088), and **d** the number of enhancers predicted by RFECS with the threshold of 0.25 in the GM12878 cell line (49699). Inside every region or intersection, the number of enhancers in the given set is indicated together with the percentage of validated enhancers in the set. The areas in the intersection sets are not proportional to the number of overlapping regions due to the asymmetry of the overlaps. . . . . . . . . . . . . . . . . . . 12

### List of Tables

S1 The classification performance (AUC) of PREPRINT and RFECS in the 5-fold CV data set from the K562 cell line and the test data from the GM12878 cell line. The methods were trained on data containing either the pure random regions or the random regions with a signal. For RFECS, the AUC values were not computed on the K562 CV data. The method with the best performance on data from different cell lines was indicated with the bold font. . . . . . . 7
S2 The number of genome-wide enhancers predicted by different methods and thresholds. . . . . . . . . . . . . . . . . . . . . . . . . . . . . . . . . . . 7

### Supplementary Methods

This sections shortly describes the five modules of the PREPRINT procedure illustrated in Figure 7. The input data for PREPRINT were the chromatin feature data obtained, e.g. by the ChIP-seq technique. In the Chromatin Immunoprecipitation (ChIP) experiment for a population of cells, the protein structures of chromatin were first covalently attached to the DNA, and the DNA was cut into small fragments, for example, by sonication. Sonication could not cut the DNA protected by the protein structures, for example, the DNA wrapped around the nucleosomes with a histone modification H3K4me1 illustrated in Figure 7**a**. Secondly, the DNA fragments attached to the chromatin feature of interest, e.g., H3K4me1, were enriched in the cell sample with a feature specific antibody. Lastly, the fragments enriched by ChIP were sequenced to produce short reads (black H3K4me1 reads in Figure 7**a**). Moreover, a set of control sequence reads were produced; these were reads obtained from the same cell population without the ChIP step to provide an estimate of the background coverage signal (black control reads in Figure 7**a**).

In the preprocessing module depicted in Figure 7**a**, the ChIP-seq and control reads were aligned back to the human reference genome. As an example, Figure 7**a** illustrates reads aligning to a 20 kb genomic region in the first human chromosome. The aligned ChIP-seq and Control reads were processed to obtain the genome-wide read coverage signal, indicated in red for the chromatin feature H3K4me1 in Figure 7**a**. The enrichment of the ChIP-seq reads and hence the high coverage signal intensity coincided with the chromosomal location of the chromatin feature. The ChIP-seq and related techniques can quantify multiple different chromatin features for the same cell line. In Figure 7**a** and in the following sections, the number of chromatin features was denoted as $K$. The number of chromatin features in the ENCODE data for cell lines K562 and GM12878 was 15.

In the second step of the preprocessing module (Figure 7**b**), the coverage signals were extracted at the training data samples to build the training data coverage matrices. The training data samples included enhancers (Class 1), promoters (Class 0), and random genomic locations (Class 0). For more information on the definition of the training data samples, see the Section Definition of the training data. For example, the training data enhancers were coordinates to single base pairs in the genome. These coordinates are referred to as anchor points, and they are indicated as the dashed line and the coordinate 0 in Figure 7**b**. Centred at the anchor point, a genomic window of size 2 kb was defined, and the window was divided into 100 bp bins. Adopting the 100 bp bin size indicates that the resolution of the coverage signal was 100 bp. For every bin along the windows, the coverage signals were extracted, resulting in a matrix of size $n \times d$. Here, $n$ is the number of the training data enhancer samples, and $d$ is the length of the row vector. With the genomic window of 2 kb and the bin size of 100, the length $d$ is 20. The row vector of the matrix is referred to as a feature pattern vector for a given sample, and the column-wise average signal of the matrix is referred to as an aggregate pattern of, for example, the enhancer class. The feature patterns in the matrix were visualised as a heatmap together with the aggregate pattern to reveal the biological properties of enhancers. The visualisations were generated by the functions in the EnrichedHeatmap bioconductor package [1]. The leftmost heatmap and the aggregate pattern in Figure 7**b** represent the coverage matrix of the chromatin feature H3K4me1 at the training data enhancers. The heatmaps and aggregate profiles for all 15 chromatin features at the training enhancers are illustrated in Figure 8.

The PREPRINT classifier is based on supervised learning. The classifier should learn to distinguish the enhancers (Class 1) from the non-enhancer regions (Class 0). As examples of non-enhancer class, promoters and random genomic regions were utilised. For details on defining the training data promoters and random genomic locations, see the Section Definition of the training data. The coverage matrices of H3K4me1 at the training data promoters and random regions are again illustrated as heatmaps together with the aggregate patterns in Figure 7**b**. The heatmaps and aggregate patterns for all 15 chromatin features at the training promoters and random regions are illustrated in Supplementary Figures S1, S2, and S3, Additional File 1.

According to the aggregate pattern of the chromatin feature MNase-seq at the enhancers presented in Figure 8, there were two well-positioned nucleosomes flanking the anchor points of the enhancers, and the nucleosomes were more mobile when moving further from the anchor points. Co-localising with the two well-positioned nucleosomes, several histone modifications, such as H3K4me1, formed bimodal peaks. In addition to enhancers, these characteristic aggregate patterns of different chromatin features can also be defined for the promoters and random regions (see Supplementary Figures S1, S2, and S3, Additional File 1). The aggregate patterns of the chromatin feature H3K4me1 for the enhancers, promoters and random regions are demonstrated in the statistical modelling module of PREPRINT (Figure 7**c**). The aggregate patterns are indicated as $\mathbf{x}^{\text{enh}}$, $\mathbf{x}^{\text{prom}}$, and $\mathbf{x}^{\text{rand}}$. Given the characteristic enhancer, promoter, or random aggregate pattern, scaled by the scaling factor $\alpha_i^{\text{enh}}$, $\alpha_i^{\text{prom}}$, or $\alpha_i^{\text{rand}}$, respectively, the elements of the individual samples were assumed to follow independent Poisson distributions (see the likelihood function in Figure 7**c**). In Figure 7**c**, on the left side, a training data enhancer ($\mathbf{y}_i^{\text{enh}}$) and a non-enhancer ($\mathbf{y}_i^{\text{rand}}$) are exemplified. The aggregate patterns multiplied by the scaling factor ($\alpha_{1i}\mathbf{x}^{\text{enh}}$) are depicted as black over the purple training data enhancer or the non-enhancer sample. The fit of the sample to the aggregate patterns is an essential concept of the PREPRINT procedure. To quantify the fit between the aggregate pattern and the individual sample, two probabilistic distance measures were defined: PREPRINT ML and PREPRINT Bayesian.

Computing the probabilistic distance measures or scores and building the final training data matrix were the steps of the third module of the PREPRINT procedure (Figure 7**d**). The distance measures were computed as follows. Firstly, the maximum likelihood (ML) estimates of the scaling parameters $\alpha_i$ were obtained for the training data enhancer samples. These ML estimates were employed to compute the probabilistic distance measures in the PREPRINT ML approach. Secondly, the ML estimates were utilised to fit Gamma distributions to obtain the (empirical) priors for the scaling parameters. The priors were employed to compute the probabilistic distance measure in the PREPRINT Bayesian approach. Finally, in the ML approach, the fit of the sample to the aggregate pattern was computed as a likelihood utilising the ML estimates whereas in the Bayesian approach, the fit was computed as the posterior predictive values obtained by integrating the likelihood over the

Gamma prior distribution. For each training data sample, the fit was computed between the sample and the enhancer, promoter, and random aggregate pattern. Therefore, the length of the final probabilistic score vector was 3. The probabilistic scores of PREPRINT ML or PREPRINT Bayesian were computed for all training data samples and $K$ different chromatin features to obtain a final training data matrix of size $4n \times 3K$ (Figure 7**e**). Performing the statistical modelling of the chromatin features (Figure 7**d**) and computing the probabilistic scores (Figure 7**e**) can be recognised as a means to reduce the dimensionality of the original, considerably high dimensional ChIP-seq data of multiple chromatin features.

The fourth module in the PREPRINT procedure was to employ the final training data matrix to train a support vector machine (SVM) classifier with a Gaussian kernel (Figure 7**f**). Finally, in the fifth module, the PREPRINT classifier was applied to predict enhancers along the whole human genome. The coverage signals at subsequent 2 kb sliding windows shifted by 100 bp along the human genome were extracted to compute the probabilistic scores (Figure 7**g**). The probabilistic scores were classified by PREPRINT resulting in genome-wide prediction scores (Figure 7**h**). The genomic regions with a prediction score above a chosen threshold were predicted as enhancers. To pinpoint the exact location of the enhancer, a single window with the maximal prediction score within the larger prediction region was selected. The exact locations of enhancers are depicted as blue bars in Figure 7**h**. In the Methods section, the input data, the PREPRINT modules, and the data processing and analysis steps within each module are described in more detail.

### References

[1] Gu Z, Eils R, Schlesner M, Ishaque N. EnrichedHeatmap: an R/Bioconductor package for comprehensive visualization of genomic signal associations. BMC genomics. 2018;19(1):234.

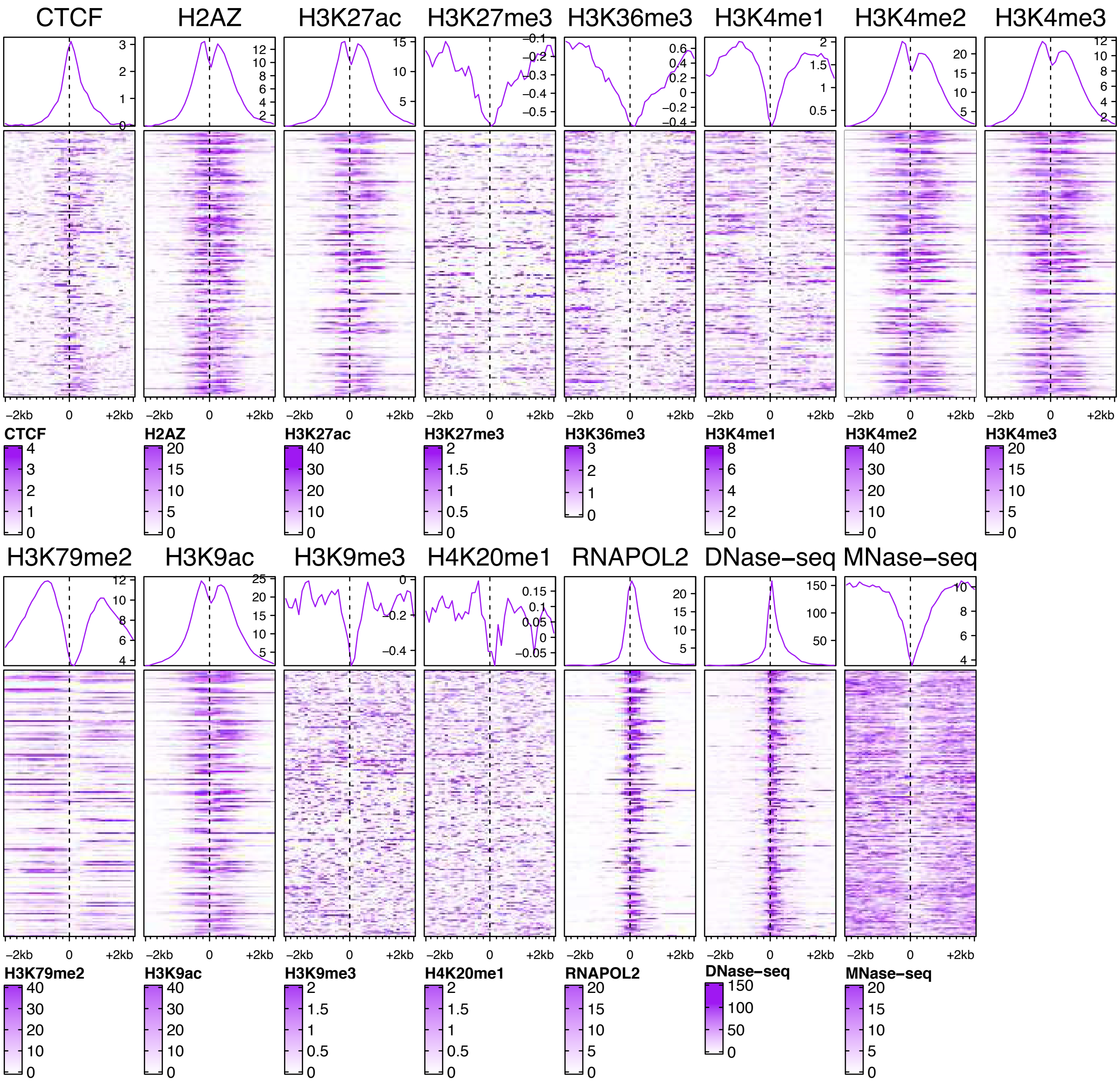


Figure S1: The coverage matrices and the aggregate patterns of the 15 chromatin features at the 1000 promoter samples. The coverage matrices were visualised as heatmaps together with the aggregate patterns illustrated above the heatmaps. The data originated from the K562 cell line, and the feature patterns were extracted in a genomic window of length 4 kb centred at the promoter anchor points (TSS) indicated by the dashed line and the coordinate 0. The resolution (bin size) of the data was 100 bp. The promoters are unoriented, i.e., the direction of transcription from TSS is not utilised to direct the coverage patterns.

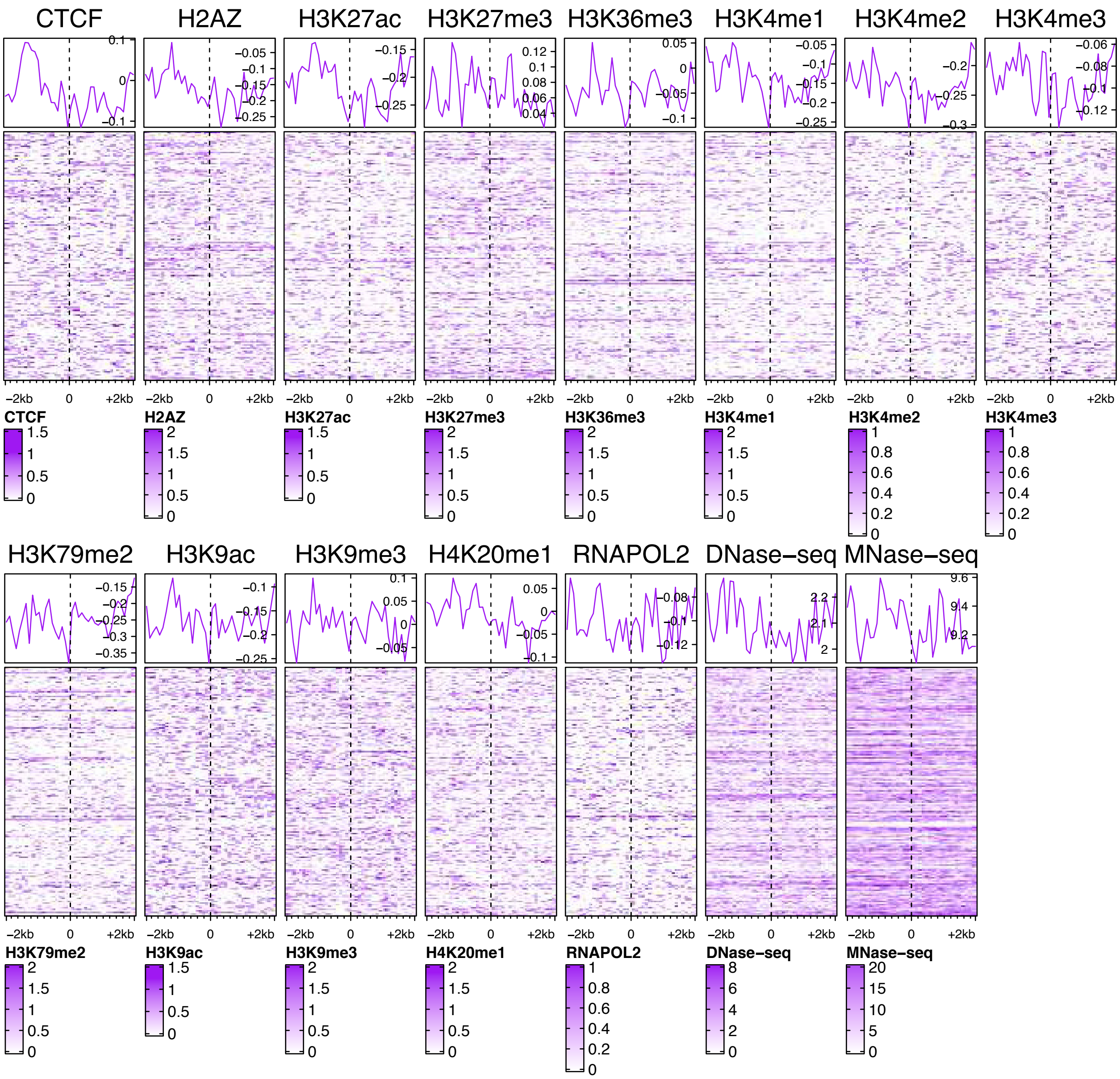


Figure S2: The coverage matrices and the aggregate patterns of the 15 chromatin features at the 1000 pure random samples. The coverage matrices were visualised as heatmaps together with the aggregate patterns illustrated above the heatmaps. The data originated from the K562 cell line, and the feature patterns were extracted in a genomic window of length 4 kb centred at the anchor points of the random regions, indicated by the dashed line and the coordinate 0. The resolution (bin size) of the data was 100 bp.

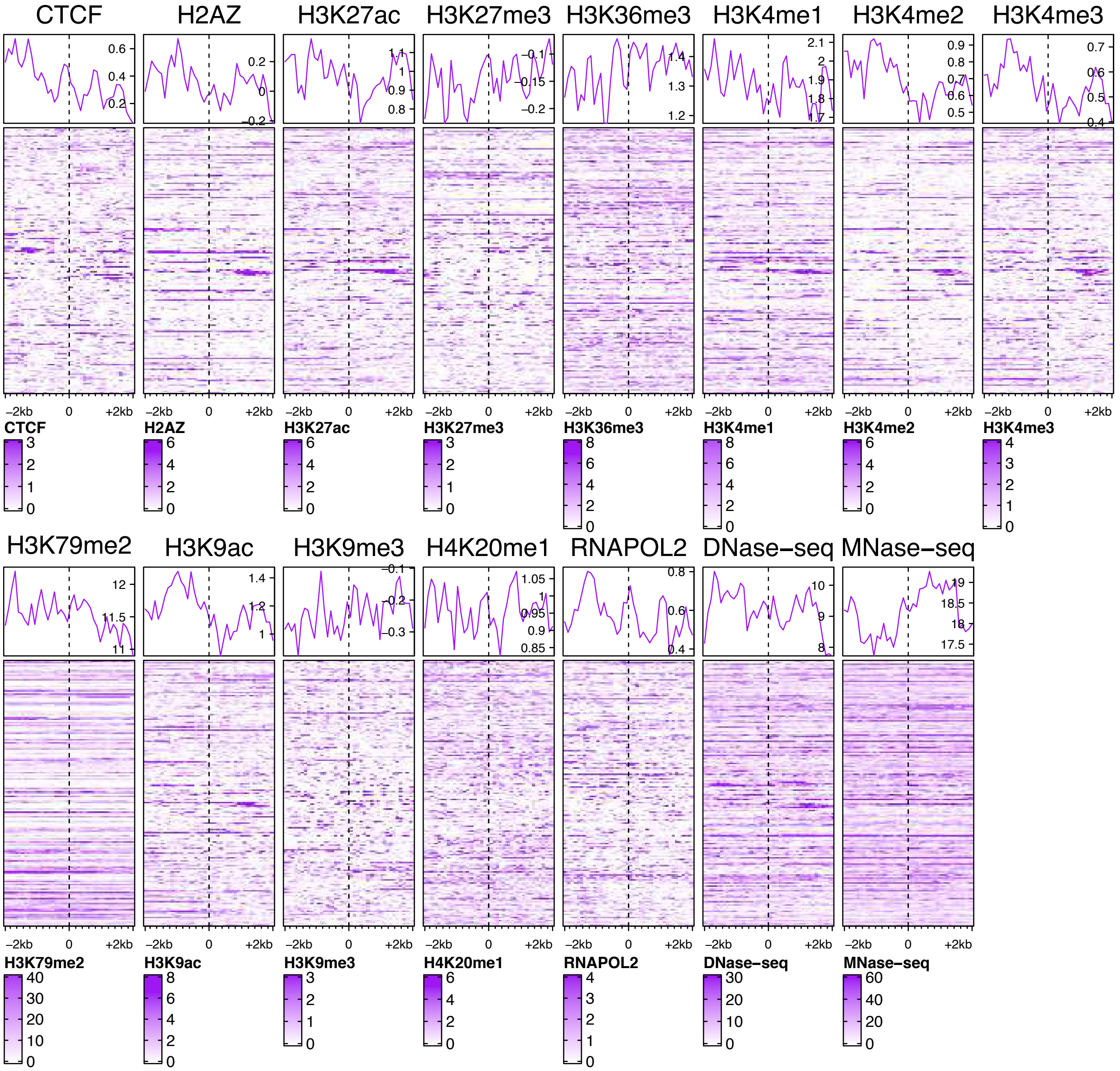

Figure S3: The coverage matrices and the aggregate patterns of the 15 chromatin features at the 1000 random regions with a signal. The coverage matrices were visualised as heatmaps together with the aggregate patterns illustrated above the heatmaps. The data originated from the K562 cell line, and the feature patterns were extracted in a genomic window of length 4 kb centred at the anchor points of the random regions, indicated by the dashed line and the coordinate 0. The resolution (bin size) of the data was 100 bp.

Table S1: The classification performance (AUC) of PREPRINT and RFECS in the 5-fold CV data set from the K562 cell line and the test data from the GM12878 cell line. The methods were trained on data containing either the pure random regions or the random regions with a signal. For RFECS, the AUC values were not computed on the K562 CV data. The method with the best performance on data from different cell lines was indicated with the bold font.

| | | AUC | |
|---|---|---|---|
| Method | Cell line | Pure random regions | Random regions with a signal |
| PREPRINT Bayesian | K562 | **0.994** | 0.989 |
| PREPRINT ML | K562 | 0.992 | **0.990** |
| PREPRINT Bayesian | GM12878 | **0.991** | **0.972** |
| PREPRINT ML | GM12878 | 0.990 | **0.972** |
| RFECS | GM12878 | 0.988 | 0.950 |

Table S2: The number of genome-wide enhancers predicted by different methods and thresholds.

| | | | The threshold of 0.5 | | The best operating point or the threshold of 0.25 for RFECS | | | The FPR 1% Threshold | | |
|---|---|---|---|---|---|---|---|---|---|---|
| Method | Cell | random | all | without TSS | threshold | all | without TSS | threshold | all | without TSS |
| PREPRINT ML | K562 | pure | 94957 | 79109 | 0.57686 | 86783 | 72241 | 0.67285 | 76660 | 63839 |
| PREPRINT Bayesian | K562 | pure | 75410 | 60338 | 0.55273 | 70795 | 56675 | 0.69234 | 59087 | 47291 |
| RFECS | K562 | pure | 31331 | 25155 | 0.25 | 65515 | 51743 | | | |
| PREPRINT ML | K562 | with a signal | 51030 | 42657 | 0.54916 | 44750 | 37188 | 0.7141 | 28947 | 23691 |
| PREPRINT Bayesian | K562 | with a signal | 42844 | 35622 | 0.51043 | 41677 | 34634 | 0.78752 | 20184 | 16467 |
| RFECS | K562 | with a signal | 18960 | 15790 | 0.25 | 44536 | 35102 | | | |
| ChromHMM Weak Enhancer | K562 | | 180471 | 176912 | | | | | | |
| ChromHMM Strong Enhancer | K562 | | 69019 | 66888 | | | | | | |
| PREPRINT ML | GM12878 | pure | 102871 | 89101 | 0.57686 | 93094 | 80818 | 0.67285 | 81113 | 70671 |
| PREPRINT Bayesian | GM12878 | pure | 77304 | 66978 | 0.55273 | 72817 | 63200 | 0.69234 | 60830 | 53251 |
| RFECS | GM12878 | pure | 34619 | 30407 | 0.25 | 99013 | 86056 | | | |
| PREPRINT ML | GM12878 | with a signal | 49679 | 44651 | 0.54916 | 43975 | 39577 | 0.7141 | 28632 | 25915 |
| PREPRINT Bayesian | GM12878 | with a signal | 44684 | 40127 | 0.51043 | 43682 | 39219 | 0.78752 | 20605 | 18702 |
| RFECS | GM12878 | with a signal | 20071 | 17758 | 0.25 | 58600 | 49739 | | | |
| ChromHMM Weak Enhancer | GM12878 | | 178474 | 175487 | | | | | | |
| ChromHMM Strong Enhancer | GM12878 | | 64052 | 62599 | | | | | | |
| The thresholds estimated for the GM12878 cell line | | | | | | | | | | |
| PREPRINT ML | GM12878 | pure | | | 0.1753 | 169042 | 147212 | 0.83915 | 59170 | 52063 |
| PREPRINT Bayesian | GM12878 | pure | | | 0.16546 | 122545 | 105304 | 0.8303 | 47060 | 41642 |
| PREPRINT ML | GM12878 | with a signal | | | 0.23674 | 103059 | 93116 | 0.82949 | 18906 | 17190 |
| PREPRINT Bayesian | GM12878 | with a signal | | | 0.25712 | 84806 | 76480 | 0.83575 | 17059 | 15527 |

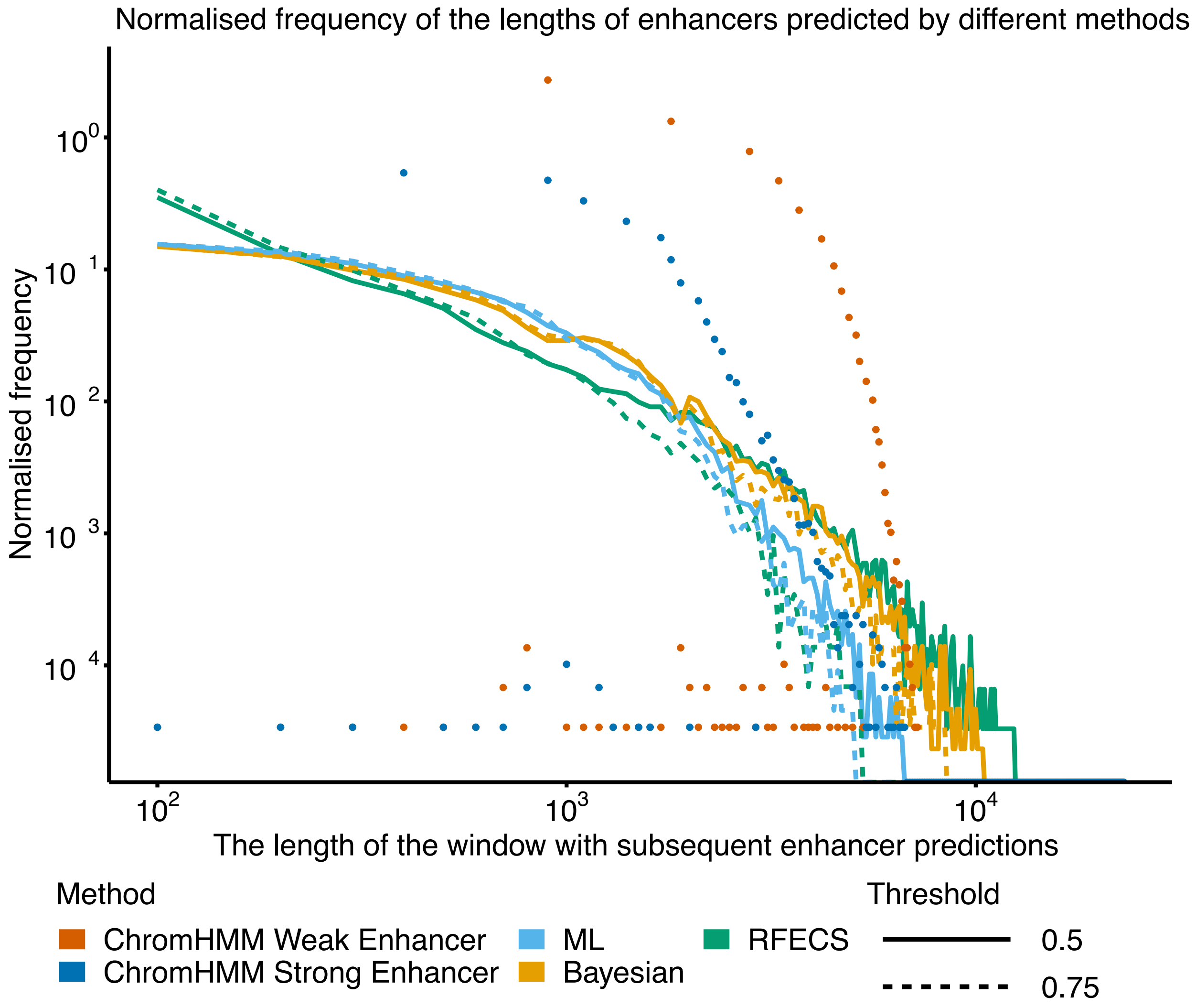


Figure S4: The normalised frequencies of the enhancer lengths. The enhancers were predicted in the GM12878 cell line by PREPRINT and RFECS with the thresholds of 0.5 and 0.75. For each method and threshold, the frequencies were divided by the total number of regions predicted as enhancers. The regions were formed by combining the subsequent enhancer predictions into a single region.

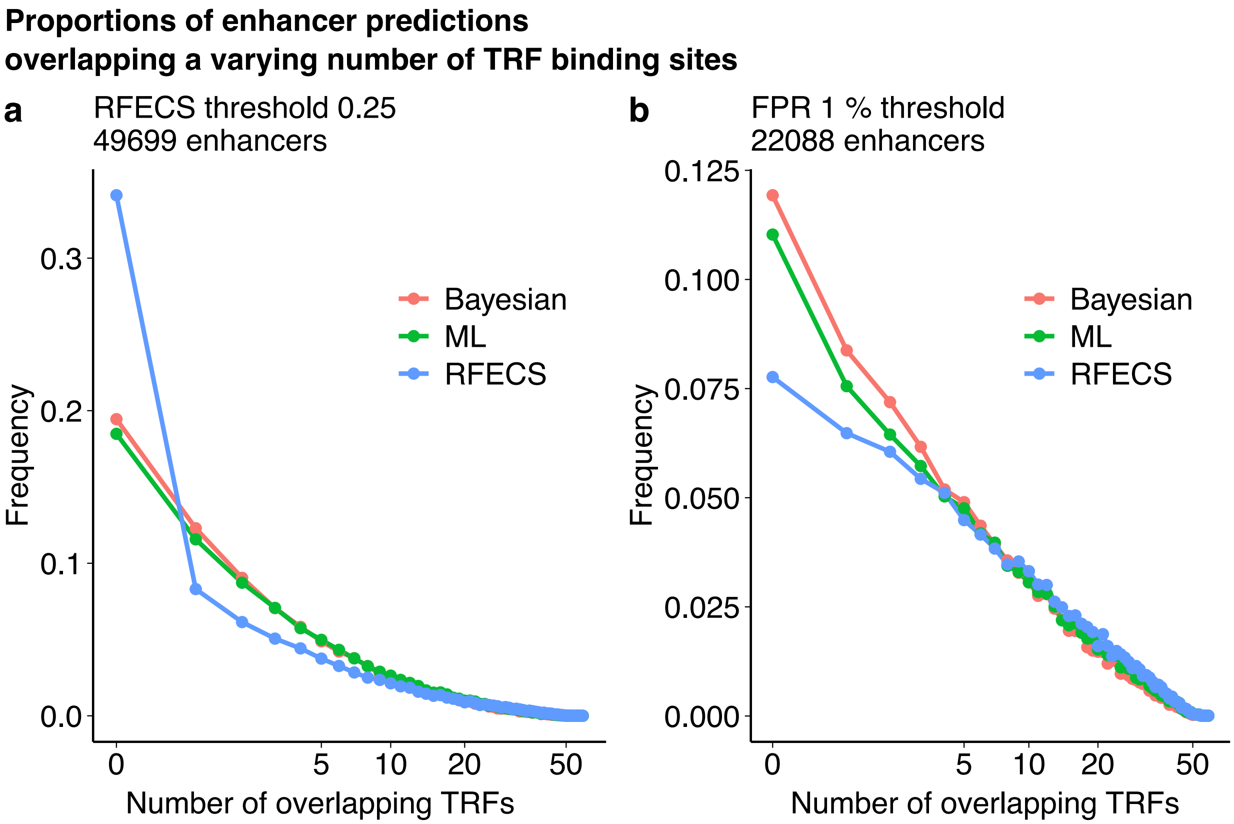


Figure S5: The proportions of the genome-wide enhancer predictions overlapping the varying number of TRF ChIP-seq peaks in the GM12878 cell line. The number of enhancers in each comparison are shown above the figure. In comparison **a**, the number of enhancers was the number of enhancers predicted by RFECS with the threshold of 0.25, and in comparison **b**, the number of enhancers was the minimum number of enhancers predicted by PREPRINT methods with their 1% FPR thresholds estimated on the K562 CV data.

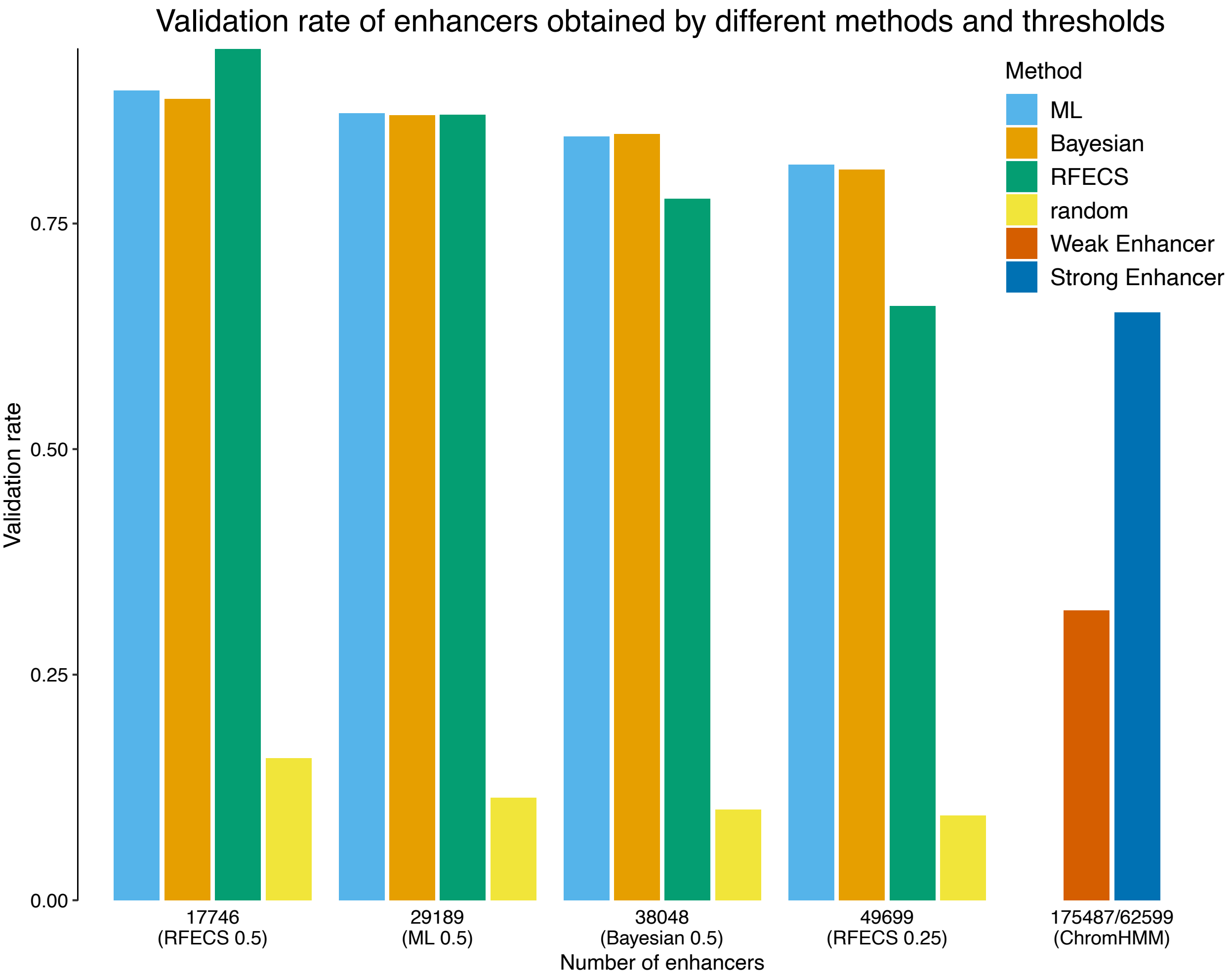


Figure S6: The validation rate of the genome-wide enhancer predictions obtained by the different methods and thresholds in the GM12878 cell line. An enhancer prediction was validated if the 2 kb prediction window overlapped with at least 1 bp of at least one TRF ChIP-seq peak.

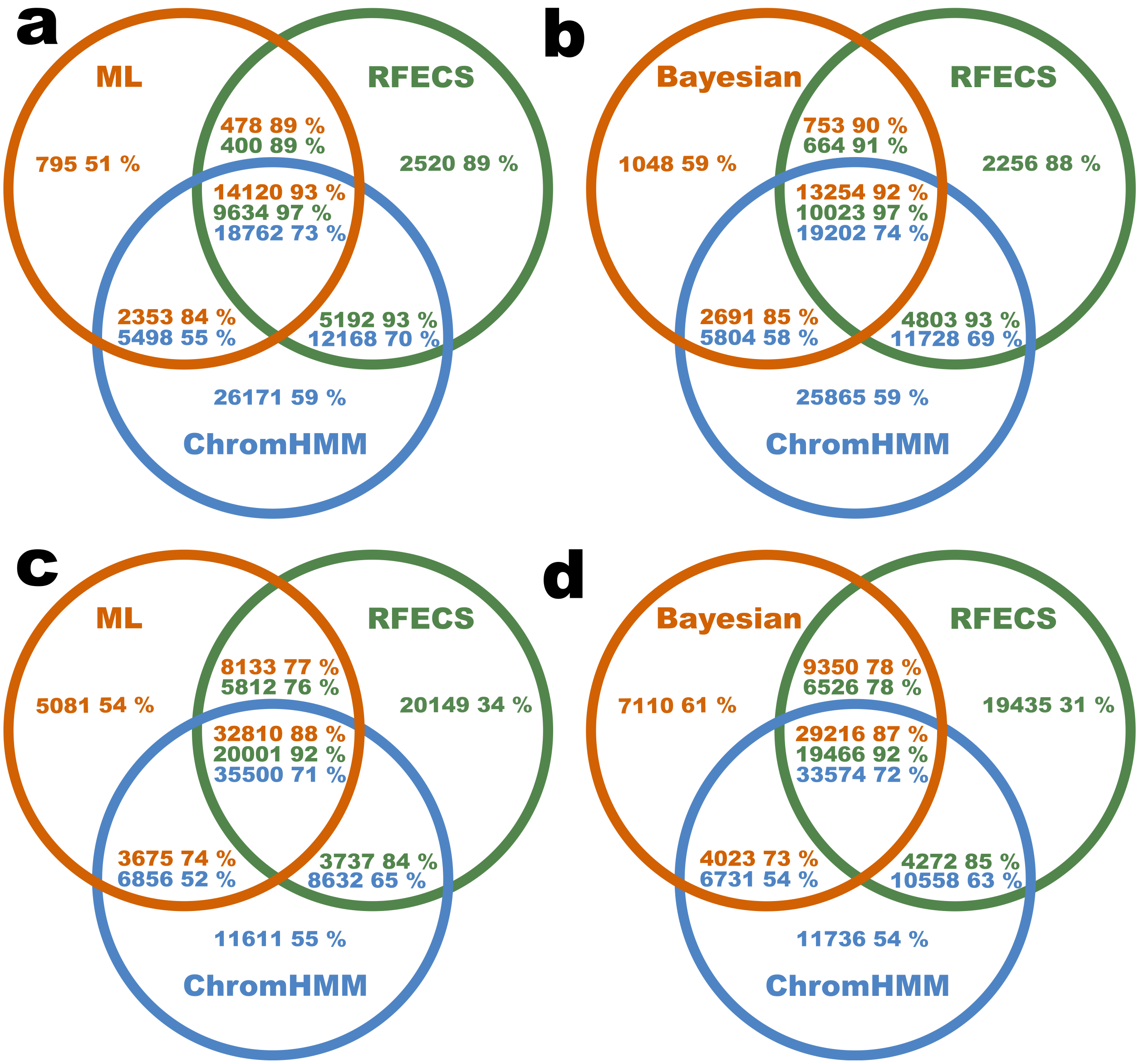


Figure S7: The unique and overlapping genome-wide enhancer predictions obtained by different methods in the GM12878 cell line. In comparisons **a** and **c**, the PREPRINT predictions were obtained by the ML approach, and in comparisons **b** and **d**, the predictions were obtained by the Bayesian approach. The overlap between the PREPRINT, RFECS and ChromHMM predictions were quantified as the number of enhancers. In each comparison, the number of enhancers predicted by PREPRINT and RFECS was equal. The numbers were: **a** 17746, **b** 17746, **c** 49699, and **d** 49699. Inside every region or intersection, the number of enhancers in the given set is indicated together with the percentage of validated enhancers in the set. The areas of the intersection sets are not proportional to the number of overlapping regions due to the asymmetry of overlaps.

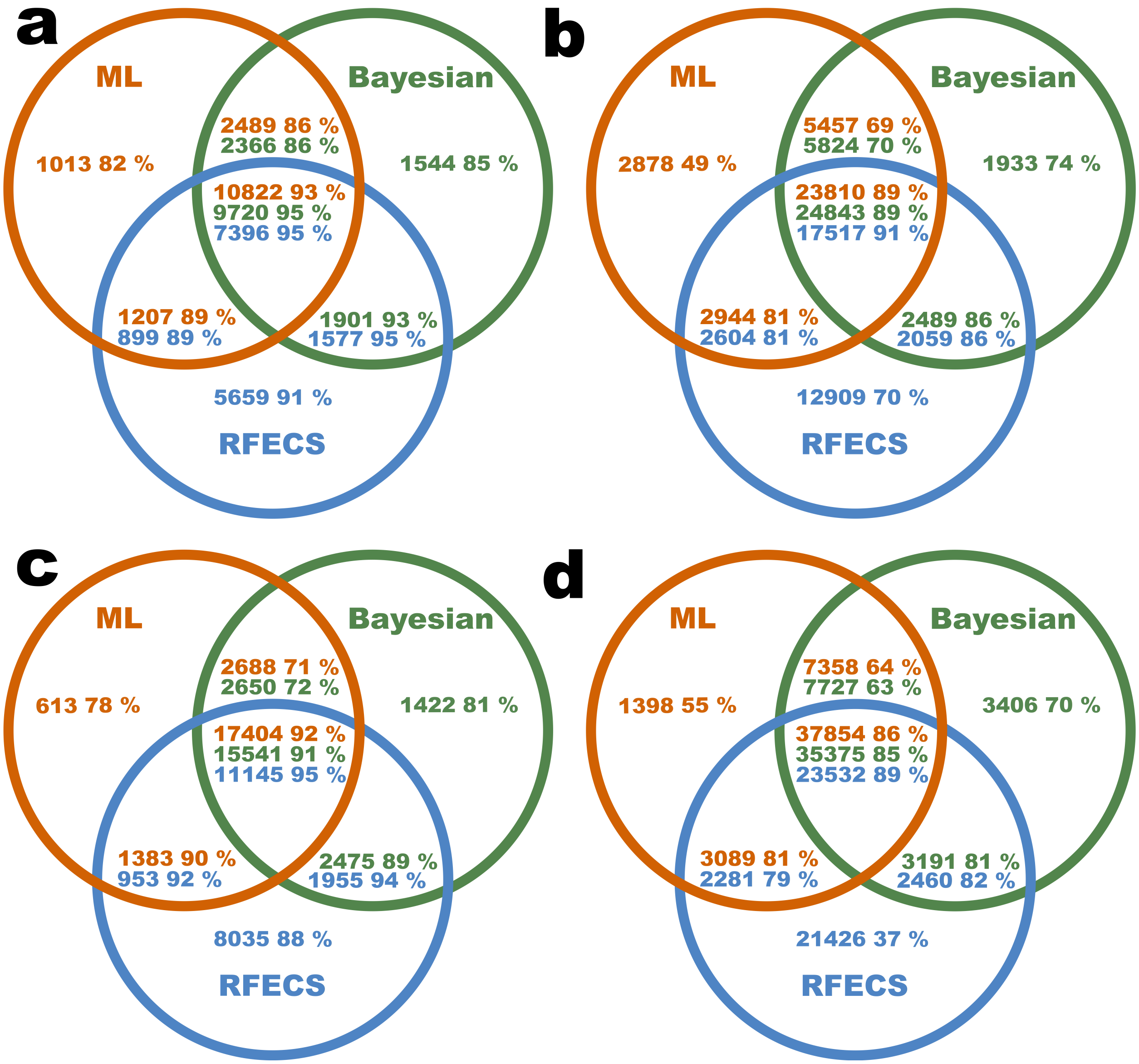


Figure S8: The unique and overlapping genome-wide enhancer predictions obtained by PREPRINT and RFECS in the K562 cell line (**a** and **b**) and in the GM12878 cell line (**c** and **d**). In each comparison, the number of enhancers predicted by PREPRINT and RFECS was equal. The numbers were: **a** the minimum number of enhancers predicted by PREPRINT with the 1 % FPR threshold in the K562 cell line (15531), **b** the number of enhancers predicted by RFECS with the threshold of 0.25 in the K562 cell line (35089), **c** the minimum number of enhancers predicted by PREPRINT with the 1 % FPR threshold in the GM12878 cell line (22088), and **d** the number of enhancers predicted by RFECS with the threshold of 0.25 in the GM12878 cell line (49699). Inside every region or intersection, the number of enhancers in the given set is indicated together with the percentage of validated enhancers in the set. The areas in the intersection sets are not proportional to the number of overlapping regions due to the asymmetry of the overlaps.
